# Bio-Templated Zinc Oxide Nanotherapeutics from *Flemingia macrophylla* kuntze Ex Merr: *In Vivo* Anti-Diabetic and Anti-Inflammatory Efficacy in *Drosophila melanogaster*

**DOI:** 10.64898/2026.10.20.760481

**Authors:** Lhaineichong Khongsai, Rounab Sarkar, Atom Rajiv Singh, Asem Sanjit Singh, Birendra Kumar Bindhani, N Upendra, Ch. Brajakishor Singh

## Abstract

Nanotechnology has a major impact on the creation of novel and effective therapies for the treatment of several diseases, including diabetes and cancer. The current work is a report on the use of the aqueous extract from the leaves of the *Flemingia macrophylla* plant as a capping and reducing agent in the green synthesis of zinc oxide nanoparticles (ZnO-NPs).The synthesized ZnO-NPs were subjected to characterization through X-ray diffraction (XRD), Fourier transform infrared spectroscopy (FT-IR), transmission electron microscopy (TEM) scanning electron microscopy (SEM), and ultraviolet-visible spectroscopy (UV-Vis). Dose-dependent *in vivo* effects were investigated using *Drosophila melanogaster* subjected to a high-sugar diet (HSD) to induce a diabetic-like model. *In vitro* anti-oxidant, anti-diabetic, anti-inflammatory assay, survival, locomotor function, neuroanatomical integrity, ovary and testis morphology, egg-laying capacity, and male fertility were evaluated. The most prominent UV-visible peak was detected at around 370 nm. Structural and morphological analysis via XRD and electron microscopy confirmed the hexagonal phase of the ZnO-NPs, indicating an average calcined particle size of around 47 nm at 500°C.The manuscript reports significant anti-diabetic and anti-inflammatory effects in vitro and in the *in vivo Drosophila* model, together with antioxidant activity. The *in vivo* results indicate that green-synthesized ZnO-NPs mitigate adverse phenotypes triggered by a high-sugar diet and also potential as an anti-diabetic intervention and warrant further mechanistic and preclinical investigation.

## 1. Introduction

Diabetes mellitus (DM) is a serious metabolic disease that occurs as a result of elevated blood glucose, oxidative stress, and chronic low-grade inflammation playing an important role in the development of insulin dysfunctions in DM patients and the appearance of the complications they experience

(Badeggi et al., 2020) Although current anti-diabetic therapies primarily target glycaemic control, their ability to simultaneously address diabetes-associated oxidative and inflammatory processes remains limited. Therefore, the development of multifunctional therapeutic strategies capable of modulating both glucose metabolism and inflammation are of considerable interest.

Nanotechnology offers new opportunities for improving the biological efficacy of therapeutic agents through the unique physicochemical properties of nanoparticles. Among these, zinc oxide nanoparticles (ZnO-NPs) have attracted considerable attention because of their nanoscale properties and the biological importance of zinc in glucose metabolism, insulin signalling, oxidative homeostasis, and cellular regulation (Issam et al.,2021;Jiang et al.,2018;Tang,2019) Increasing evidence suggests that ZnO-NPs possess antidiabetic and anti-inflammatory potential, including effects on glucose regulation and inflammatory signalling pathways such as NF-κB (Thema et al.,2015; Mishra et al.,2017; Paul et al.,2020; Aamri et al.,2024; Ikbal et al.,2023)These properties make ZnO-NPs promising candidates for targeting the interconnected metabolic and inflammatory abnormalities associated with diabetes.

Green synthesis using medicinal plant extracts provides an environmentally sustainable approach for producing ZnO-NPs while utilizing naturally occurring phytochemicals as reducing, capping, and stabilizing agents (Sharma et al.,2023; Aldeen et al.,2022; Govindan et al., 2020). Importantly, plant-derived flavonoids, phenolics, alkaloids, and other bioactive compounds may contribute additional antioxidant and anti-inflammatory activities to the resulting nanoparticles (Ochieng et al.,2015). Thus, plant-mediated ZnO-NPs may provide a multifunctional nanoplatform in which the biological properties of ZnO are complemented by the pharmacological activities of phytochemicals.

*Flemingia macrophylla* (Willd.) kuntze ex Merr. (Fabaceae) is a medicinal plant containing bioactive compounds such as kaempferol, rutin and quercetin which have reported antioxidant and anti-inflammatory properties (Gahlot et al.,2013). Other phytochemical investigations have also identified biologically active alkaloidal constituents (Kumbhare & Ukey 2023; Singleton et al.,1999; Marinova et al.,2005). However, the antidiabetic and anti-inflammatory potential of ZnO-NPs synthesized using *Flemingia macrophylla (F. macrophylla*) has not been adequately investigated.

Therefore, the present study aimed to synthesize and characterize ZnO-NPs using an aqueous leaf extract of *F. macrophylla* and systematically evaluate their antidiabetic and anti-inflammatory efficacy using in vitro and *in-vivo Drosophila melanogaster*-based models. The study further examined their dose-dependent biological safety to establish the relationship between therapeutic efficacy and toxicity. This integrated approach provides a basis for evaluating *F. macrophylla*-mediated ZnO-NPs as a potential multifunctional nano biological strategy for diabetes and its associated inflammatory complications.

## 2. Material and Methods

### 2.1. Plant material collection, identification and voucher specimen

*F. macrophylla fresh* leaves were collected from the hill range of Andro village, Imphal, Manipur (Latitude 24°43’42.33’’N, Longitude 94°03’37.56’’E and Altitude 815m above sea level). The plant identification was done by plant taxonomist at Botanical Survey of India (BSI), Eastern Regional Centre, Shillong, Meghalaya. The botanical specimen of the identified plant species was deposited at the depository library BSI Shillong (Voucher number: BSI/ERC/Tech/2021-22/528)

### 2.2. Preparation of aqueous plant extract

The freshly plucked leaves of *F. macrophylla* were washed with distilled water for 4-5 times to remove all the impurities (dust) present in it, and then were finely chopped. 30g of finely chopped leaves were soaked in 250mL distilled water in a conical flask and heated at 60°C for 45min.The yellowish color extract from the plant was cooled and filtered with the help of Whatman No. 1(filter paper) to remove all the remains, and the filtrate was stored in a refrigerator at 4°C for further analysis (Electronic spectra and biological applications).

### 2.3. Biosynthesis of green ZnO-NPs

To synthesize ZnO-NPs for the experiments, a slight modification of the green synthesis technique from a previous study was used (Naseer et al.,2020) Zn(CH_3_COO)_2_.2H_2_O (25 mmol, 4.587 g) was dissolved in 100 mL distilled water with continuous stirring. 20 mL of aqueous plant extract and 10mL of 1M NaOH were added to the zinc acetate solution under stirring (600 ppm) at 60°C for 6h. The mixture sample was then cooled and kept at room temperature for 24 hours to settle the precipitate formed. The clear solution was decanted, and the remaining solution was centrifuged at 10000 rpm for 10 min with continuous washing with distilled water and absolute ethanol for three times so as to remove all the impurities. The precipitate collected was dried at 90°C at oven for 24h and then calcinated at 500°C for 2h in a muffler furnace. The collected green ZnO-NPs were used for characterization and biological application studies.

**Scheme 1.**
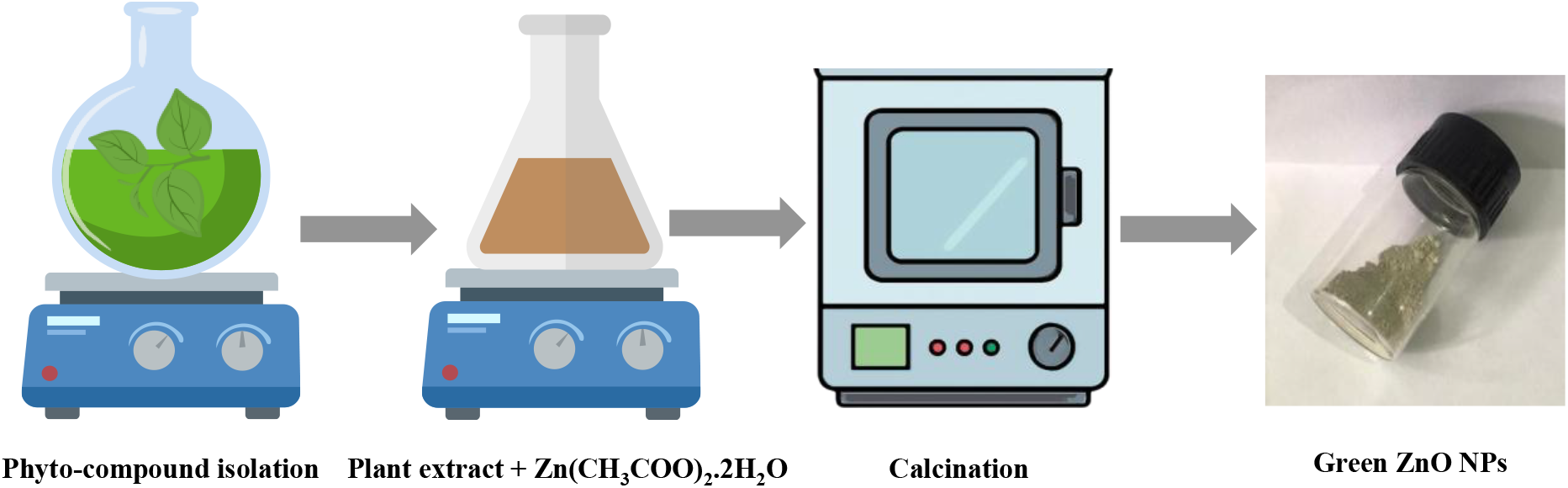
The stepwise biogenic fabrication of ZnO-NPs from precursor solution to calcined crystals using *Flemingia macrophylla* extract

### 2.4. Characterization Methods

FT-IR spectra of the prepared green ZnO-NPs were measured with JASCO FT-IR 4100 spectrophotometer using KBr pellets in the range of 4000-400 cm^-1^. Powder X-ray diffraction (PXRD) of the crystal structure of the prepared green ZnO-NPs (annealed at 500°C) was recorded using Phillip’s PANAlytical X’PERT PRO Diffracto meter with Cu target (λ=1.5404) in 2θdegrees from 10-80ranges. Electronic spectra of green ZnO-NPs were recorded using Shimadzu 1601 (Tokyo, Japan) UV-Vis spectrophotometer. TEM images and selected area electron diffraction (SAED) patterns were recorded in a Jeol2100 transmission electron microscope with field emission gun operating at 200 kV.

### 2.5. Determination of biological potential of the green synthesized ZnO-NPs

#### 2.5.1. Preliminary phytochemical characterization

##### 2.5.1.1 Total phenolic content

Total phenolic content (TPC) was determined using the Folin–Ciocalteu method according to Singleton et al. (1999) with slight modifications. Briefly, 100 µL of ZnO-NPs sample (1 mg/mL) was mixed with 500 µL of 10% Folin–Ciocalteu reagent and incubated at room temperature for 10 min. Subsequently, 400 µL of 7% Na_2_CO_3_ was added, and the mixture was incubated at 37±1°C for 30 min. Absorbance was measured at 760 nm using a microplate reader. Gallic acid (10-100 µg/mL) was used to generate the calibration curve. TPC was calculated from the standard curve and expressed as mg gallic acid equivalents (GAE)/g extract. Measurements were performed in triplicate and reported as mean±SD.

##### 2.5.1.2. Total amount of flavonoids

Total flavonoid content (TFC) was determined using the aluminium chloride colorimetric method described by Marinova et al. (2005). Briefly, 100 µL of ZnO-NPs sample (1 mg/mL) was mixed with 400 µL water and 30 µL of 5% NaNO_2_ and incubated for 5 min. Subsequently, 30 µL of 10% AlCl_3_ was added and incubated for 6 min, followed by 240 µL water and 200 µL of 1M NaOH. After 15 min at room temperature, absorbance was measured at 510 nm. Catechin (1-100 µg/mL) was used to generate the calibration curve, and TFC was expressed as mg catechin equivalents (CE)/g extract.

#### 2.5.3. *In-vitro* antioxidant assay of the green synthesized ZnO-NPs

The anti-oxidant potential of the green-synthesized ZnO nanoparticles (ZnO-NPs) was evaluated using complementary free-radical scavenging assays, including the 2,2-diphenyl-1-picrylhydrazyl (DPPH) and 2,2′-azinobis-(3-ethylbenzothiazoline-6-sulfonic acid) (ABTS) assays and reducing power assay. These assays were selected to assess the ability of the synthesized nanoparticles to neutralize chemically generated free radicals through hydrogen atom or electron transfer mechanisms. As the ZnO-NPs were synthesized using F. macrophylla leaf extract, the observed anti-oxidant activity may be associated not only with the physicochemical properties of the nanoparticles but also with bioactive phytoconstituents adsorbed or incorporated during the green synthesis process

##### 2.5.1.1. DPPH radical scavenging assay

The anti-oxidative activity of the biosynthesized ZnO-NPs was determined using 1,1-diphenyl-2-picrylhydrazyl (DPPH) radical scavenging assay (Baliyan et al.,2022) DPPH stock solution was freshly prepared by dissolving 5mgof DPPH in 25 mL of methanol. The solution was then diluted with methanol to obtain absorbance values within an optical density range of 0.8 to 1.0. Further, 2 mL of the DPPH solution was mixed with 25µL each sample at concentrations of (10-100 μg/mL). The mixture was incubated for 30 minutes, and the absorbance was read at 517 nm using a spectrophotometer (Shimadzu UV-1601, Tokyo, Japan). Methanol was used as the control, and DPPH solution as the control reference, while ascorbic acid was used as the standard reference compound. A decrease in absorbance indicates an increase in free radical scavenging activity in the reaction mixture. Experimental tests were carried out in triplicate.

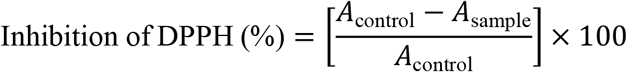

Where *A*_sample_is the absorbance of biosynthesized ZnO-NPs and *A*_control_is the absorbance of the DPPH mixture.

##### 2.5.1.2. ABTS radical scavenging assay

The ABTS (2,2′-azino-bis(3-ethylbenzothiazoline-6-sulfonicacid)) radical scavenging assay was used to investigate the anti-oxidant properties of the biosynthesized ZnO-NPs by following the method of Re et al. (1999). Approximately 20mg of ABTS was dissolved in 5 mL of methanol to prepare ABTS radical cation solution that was then mixed in a ratio of 1:1 with a solution composed of 30mg of potassium persulfate dissolved in 5 mL of methanol. The mixture was left to form the ABTS radical for 16 h at room temperature in the dark. The resulting ABTS solution was further diluted with methanol before use to give an absorbance value between 0.8 and 1.0. Each sample was added to 2 mL of the resulting ABTS solution with concentrations ranging from 10-100 μg/mL and the absorbance was read at 745 nm using a spectrophotometer after incubation for 30 minutes. All tests were carried out in triplicate.

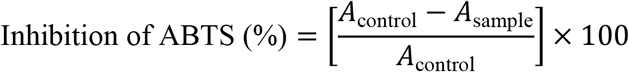

where *A*_sample_ is the absorbance of biosynthesized ZnO-NPs, and *A*_control_ is the absorbance of ABTS stock solution.

##### 2.5.1.3. Reducing power assay

The reducing assay was carried out by referring to Oyaizu et al., (1986) with slight modifications (Oyaizu,1986). Briefly, 1 mL of the test sample with concentration ranged from 10-100 μg/mL, was mixed with 2.5 mL of sodium phosphate buffer (0.2 M, pH 6.6) and 2.5 mL of potassium ferricyanide (1%). Then the reaction mixture was incubated for 20 min at 50°C water bath. After incubation, 2.5 mL of 10% trichloroacetic acid was added to the reaction mixture, then centrifuged at 3000 rpm for 10 min. Finally, 2.5 mL of the supernatant was mixed with 2.5 mL of distilled water and 0.5 mL of FeCl_3_ (0.1%) was added. The addition of 0.5 mL of FeCl_3_ (0.1%) was carried out after the supernatant was mixed with 2.5 mL of distilled water. Lastly, the absorbance (Abs) was measured at 700 nm. The increasing in an absorbance value indicates that the reaction mixture has more reducing power.

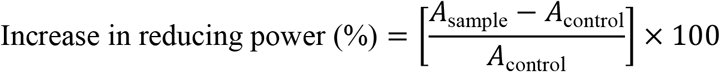

#### 2.5.2. *In vitro* assessment of the antidiabetic activity of green-synthesized ZnO-NPs through inhibition of carbohydrate- and lipid-digesting enzymes

The antidiabetic potential of green-synthesized ZnO-NPs was evaluated against the carbohydrate-hydrolyzing enzymes α-amylase and α-glucosidase, which sequentially mediate starch digestion and glucose release. Inhibition of these enzymes may reduce carbohydrate breakdown and postprandial glucose availability. Pancreatic lipase was additionally assessed as a complementary target involved in dietary triglyceride hydrolysis, where its inhibition may reduce lipid digestion and absorption.

##### 2.5.2.1. Alpha-glucosidase inhibitory assay

The *α*-glucosidase enzymatic process was performed as described by Kumar et al. (2011), with minor changes. The pre-incubation approach used p-nitrophenyl-α-D-glucopyranoside (pNPG) as a substrate. Incubate 50 μL of plant test samples at different concentrations with 25 μL of enzyme solution (0.5 U/mL in phosphate buffer) at 37±1°C for 10 min. A 25 μL solution of pNPG (0.5 mM in 0.1 M phosphate buffer, pH=6.8) was added to the reaction mixture. The mixture was incubated at 37±1°C for 30 minutes. The reaction was halted by adding 100 μL of 0.2 M Na_2_CO_3_ solution to the mixture. The absorbance was measured at 405 nm with a microplate reader (ThermoMultiskan). Acarbose served as a positive control.

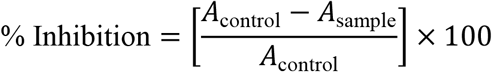

##### 2.5.2.2. α-Amylase inhibitory assay

The α-Amylase enzyme inhibitory experiment was carried out in accordance with Shu et al. (2009) In short, 50 µL of the tested solution was combined with 25 µL of the enzyme solution (0.5 mg/mL concentration) in a 0.02 M phosphate buffer solution (pH 6.9, 0.006 M NaCl) and incubated for 10 minutes at 25±1°C. After adding 50 µL of 0.5% w/v starch solution, the mixture was incubated for an additional 10 minutes at 25°C.After adding 100 µL of DNS color reagent to halt the reaction, the reaction solution was maintained in a water bath at 80 °C for ten min. The amount of residual starch was determined by measuring enzyme activity at 540 nm. An adequate blank was utilized for every sample. Acarbose was used as a standard.

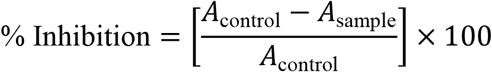

##### 2.5.2.3. Pancreatic Lipase Inhibition Assay

Pancreatic lipase inhibition was assessed using *p*-nitrophenyl butyrate (pNPB) as the substrate, following a reported method (Kim et al.,2010) with modifications. Porcine pancreatic lipase (PPL) was prepared at 1 mg/mL in potassium phosphate buffer and stored at −20°C. ZnO-NPs and orlistat (10–200 μg/mL) were pre-incubated with PPL for 60 min at 30°C in 0.1 mM potassium phosphate buffer (pH 7.2). The reaction was initiated by adding pNPB and incubated for 5 min at 30°C. Formation of *p*-nitrophenol was measured at 405 nm. PPL with pNPB without inhibitor served as the control, and lipase inhibition was expressed as a percentage of the control activity.The percentage inhibition of pancreatic lipase was calculated using the following equation:

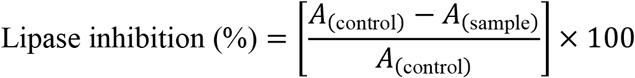

Where *A*_(control)_ represents the absorbance of the enzyme reaction in the absence of inhibitor and *A*_(sample)_ represents the absorbance obtained in the presence of the ZnO-NPs extract or orlistat. A higher percentage inhibition indicates stronger suppression of pancreatic lipase activity.

2.5.3. *In vitro* anti-inflammatory activity:

The *in vitro* anti-inflammatory potential of the green-synthesized ZnO-NPs was evaluated using protein denaturation and heat-induced hemolysis assays. These complementary models were employed as preliminary biochemical approaches to investigate the ability of ZnO-NPs to protect proteins and erythrocyte membranes against damage associated with inflammatory conditions.

##### 2.5.3.1. Bovine Serum Albumin (BSA) Denaturation Assay

The anti-inflammatory characteristics of ZnO-NPs were assessed in compliance with Das et al. (2019). The activity was measured by mixing 0.45 mL of a 1% water solution of BSA with several concentrations of ZnO-NPs extract (10 µg/mL, 50 µg/mL, 100 µg/mL, 150 µg/mL, and 200 µg/mL). To lower the pH of the solution to 6.3, an amount of 1 N HCl was added. These samples were heated to 55°C for 30 minutes in a water bath following 20 minutes at room temperature. After being heated, the samples were left to cool. At 660 nm, the absorbance was measured. The typical medication was diclofenac sodium salt. In this experiment, dimethyl sulfoxide (DMSO) was employed as a control group. Percentage of protein denaturation was estimated by the equation below;

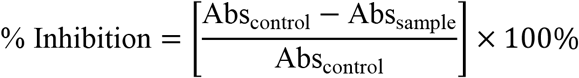

##### 2.5.3.2. Heat-induced hemolysis

A heat-induced hemolysis assay assesses the stability of red blood cell (RBC) membranes under thermal stress in order to evaluate a compound’s capacity to reduce inflammation (Chandra et al., 2012). In this experiment, an equal volume of 10% reconstituted RBC solution and different concentrations of 1 mL test samples were added to each test tube. Instead of the test sample, the control test tube contained only regular saline. The reference medication was indomethacin. Centrifuge tubes containing the reaction mixtures were incubated at 55±1°C for 30 minutes. The tubes were kept in cold water for the duration of the incubation. The absorbance at 560 nm was then measured after the tubes were spun for seven minutes at 2400 rpm. Every test sample underwent three analyzes.

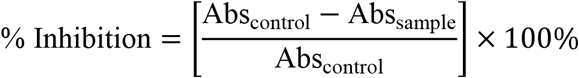

#### 2.5.4. Cell culture and MTT cell viability assay

HeLa and HEK-293T cells were cultured in DMEM supplemented with 10% heat-inactivated fetal bovine serum, sodium pyruvate (0.01 mM), NaHCO_3_ (0.2%), and penicillin-streptomycin (25 U/mL). HeLa and HEK-293T cells were utilized in ZnO-NPs cell viability experiments using the MTT assay (Mosmann,1983; Tchounwou et al.,2000). The cells were kept in a CO_2_ incubator at 37°C. Approximately 104 cells were planted in 96-well plates and incubated at 37°C for one day. Each well was then filled with varying amounts of ZnO-NPs and allowed to incubate in the dark for three to four hours. After adding MTT, the blue formazan crystals were dissolved in 100 μl DMSO (dimethyl sulfoxide). Docetaxel (25 nM) served as a positive control for the experiment. The absorbance (A) of the samples was measured at 570 nm. The percentage of cell viability was calculated using the following formula:

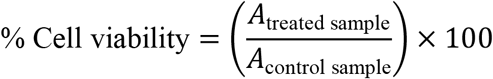

#### 2.5.6. *Drosophila melanogaster* strain and rearing conditions

The fruit fly (*Drosophila melanogaster*) is a model organism for genetic and biomedical research because of their short life cycle, similarities with humans at genetic level, and low maintenance cost. Also, around 75% of human disease-causing genes have similar genes in fruit flies which makes them an ideal candidate for studying human specific diseases like diabetes (Wangler et al.,2015; Wangler et al.,2020) Every *in vivo* experiment in this research utilized the *w^1118^*strain of *Drosophila melanogaster*. The flies were raised on a standard *Drosophila* diet based on cornmeal-sucrose-yeast agar medium, and propionic acid. The flies were kept at a temperature of 24 ± 1°C in a 12-hour light and 12-hour dark cycle, with relative humidity between 65-70%. The flies were consistently sub-cultured to sustain vibrant populations. A perishable food source was set up and changed every three days, and the flies were moved to fresh vials during this procedure. Culture vials were meticulously cleaned, rinsed, and sterilized through autoclaving prior to repurposing. This maintenance procedure was adhered to consistently during the entire period of the research (Piper et al.,2007; Bass et al.,2007)

#### 2.5.7. Statistical analysis

In order to generate a dose-response curve with various doses, all studies were conducted in triplicate and repeated three times. Data were gathered and computed as the mean standard deviation (SD) across all three distinct experiments.

GraphPad Prism 8 was used to run all statistical tests. The statistical significance of the data was assessed using the Mantel-Cox test for lifespan analysis and toxicity assessment. The fecundity, fertility, and climbing tests were all conducted in an unpaired manner. Any p-value less than 0.05 was deemed significant.

#### 2.5.8. High-sugar diet-induced diabetic-like model

The research was carefully designed to enable disease induction through dietary modifications at different concentrations, following the methods set forth by Musselman et al., 2019, which have been suitably adjusted for our study’s aims.

DM was consistently triggered in *w^1118^*strain of *D. melanogaster*by incorporating 2.5g of sucrose into every 10g of the standard diet; the remaining components of the typical fly food were kept uniform. After a span of 10 days, the flies were assessed for signs of diabetes, which included lower egg output and a decrease in movement activity (Omale et al.,2021).

#### 2.5.9. ZnO-NPs and standard metformin exposure and dose selection

Four distinct concentrations (0.5, 1.0, 2.0, and 4mg/10g diet) of ZnO-NPs extract and standard metformin (16mg/10g diet) were employed in the research. The concentrations of the extracts and standard drug were chosen following an initial toxicity and bioactivity evaluation. The extracts were incorporated into the *Drosophila* diet to evaluate toxicity in adult flies after 10 days of HSD. The control group was kept under identical conditions for all experiments without any treatment involving the extract.

### 2.6. Preparation of sample for biochemical assays

#### 2.6.1. Extraction of fly homogenate

For *in vivo* anti-oxidant and anti-diabetic assays, twenty flies each from the ZnO-NPs treated of all concentration HSD and normal flies were taken and frozen on the ice. After that, the head and body were separated, discarding the heads. The fly samples were washed thrice with 1 ml of 50 mM potassium phosphate buffer and the bodies were homogenized in HEPES buffer (20 mM, pH 7.0) as per each protocol analysis. The supernatant was collected through centrifugation at 17857 rpm for 10 minutes at 4°C and used for the biochemical experiments. All experiments were done in triplicate.

### 2.7. Estimation of glucose concentration in drosophila

Glucose concentration in the samples was determined using a glucose oxidase (GO) (Sigma-Aldrich, GAGO-20) and o-toluidine colorimetric assay kit according to the manufacturer’s protocol.

#### 2.7.1. Glucose oxidase (GO) based colorimetric assay

Glucose levels were estimated using a glucose oxidase (GO)-based colorimetric assay. Briefly, 100 µL of GO reagent was added to each sample and incubated at 37 °C for 60 min. The reaction was terminated by adding 100 µL of 12 N H_2_SO_4_, and absorbance was measured at 540 nm using a microplate reader. The resulting pink color indicated the completion of the reaction.

#### 2.7.2. o-toluidine colorimetric method

Glucose content in *Drosophila melanogaster* was estimated using the *o*-toluidine colorimetric method described by Dubowski (1962) with slight modifications. Flies were collected after treatment, rinsed with distilled water, blotted, weighed, and homogenized in ice-cold 0.1 M phosphate buffer (pH 7.0). Homogenates were centrifuged at 4,000×*g* for 10 min at 4°C, and the supernatants were used for glucose estimation. Absorbance was measured at 630 nm against a reagent blank, and glucose concentrations were determined using a standard calibration curve.

### 2.8. Lifespan Analysis

Newly eclosed male and female *D. melanogaster* (within 24 h of eclosion) were collected and maintained in groups of 40 flies per vial under normal diet (ND) and high-sucrose diet (HSD) conditions. After establishment of the HSD-induced diabetic phenotype, flies were transferred to vials containing HSD supplemented with the respective concentrations of *F. macrophylla* extract, as described above. Flies maintained on ND and HSD without extract supplementation served as the respective control groups.

For lifespan assessment, the number of dead flies in each vial was recorded at 2days interval throughout the experimental period. Following each assessment, surviving flies were gently transferred to fresh vials containing the corresponding diet and treatment to maintain consistent nutritional and treatment conditions. Mortality was monitored until the completion of the experimental period, and survival data were used to construct lifespan curves (Lee et al.,2008)

### 2.9. *In Vivo* Anti-oxidant, Anti-Diabetic and Anti-Inflammatory Efficacy in *Drosophila melanogaster*

#### 2.9.1. DPPH Radical-Scavenging Assay

The antioxidant capacity of the *D.melanogaster* samples was evaluated using the DPPH (2,2-diphenyl-1-picrylhydrazyl) radical-scavenging assay with minor modifications of the previously reported method (Johnmark & Kinyi, 2021). Briefly, 180 μL of the prepared fly supernatant was combined with 20 μL of freshly prepared 1 mM DPPH solution in methanol. The reaction mixtures were incubated for 30 min at room temperature in the dark to prevent light-induced degradation of the radical. Absorbance was subsequently recorded at 517 nm using an Infinite^®^ M Nano ELISA reader (TECAN). Each sample was analyzed in three independent measurements.

The percentage of DPPH radical-scavenging activity was calculated using:

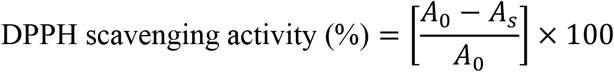

Where *A₀*represents the absorbance of the DPPH reaction mixture without fly supernatant and *Aₛ* represents the absorbance measured in the presence of the sample. To account for differences in sample protein concentration, the resulting scavenging activity was normalized to the total protein content of the corresponding supernatant.

#### 2.9.2. ABTS Radical-Scavenging Assay

The anti-oxidant activity of the *Drosophila* samples was further assessed using the ABTS [2,2′-azinobis-(3-ethylbenzothiazoline-6-sulfonic acid)] radical-scavenging assay according to the previously described procedure (Beghelli et al., 2022). Briefly, 20 μL of each fly supernatant was mixed with 180 μL of the prepared ABTS radical solution. The reaction mixtures were maintained at room temperature in the dark for 10 min, after which absorbance was measured at 734 nm using an Infinite^®^ M Nano ELISA reader (TECAN). Each sample was analyzed in triplicate.

ABTS radical-scavenging activity was determined according to the following equation:

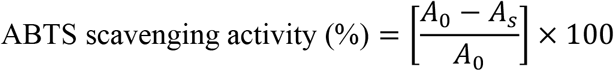

Where *A₀* denotes the absorbance of the ABTS radical solution in the absence of sample and *Aₛ* denotes the absorbance obtained following addition of the fly supernatant. The calculated antioxidant activity was subsequently normalized against the total protein concentration of each supernatant to minimize the influence of variation in protein content among samples.

#### 2.9.3. *In Vivo* Anti- Glucosidase Activity

α-Glucosidase activity in *D. melanogaster* was determined using a colorimetric assay based on the hydrolysis of *p*-nitrophenyl-α-D-glucopyranoside (pNPG), following a reported method (Wongchum et al.,2023) with modifications. Briefly, 50 µL of fly homogenate supernatant was mixed with 50 µL of 50 mM potassium phosphate buffer and 50 µL of 5 mM pNPG, followed by incubation at room temperature for 30 min. The release of *p*-nitrophenol was measured at 405 nm using an ELISA reader. Enzyme activity was calculated from a standard curve, and α-glucosidase inhibition was determined:

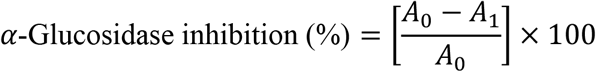

Where *A₀*is the absorbance of the untreated control and *A₁* is the absorbance of the treated sample. All measurements were performed in triplicate.

#### 2.9.4. Locomotor activity: Rapid Iterative Negative Geotaxis assay

Locomotor performance was assessed using the Rapid Iterative Negative Geotaxis (RING) assay after 15 days of dietary exposure to ZnO-NPs. Briefly, 10 flies from each treatment group were transferred to an empty vertical vial and stimulated by tapping three times. The climbing response was recorded over 10 consecutive trials, with a 120-s recovery interval between trials. Locomotor activity was compared among treatment groups based on climbing performance.

The number of flies that reached the 6 cm point of the column in 6 seconds, along with those that stayed below this mark after this duration, was recorded as described earlier (Ali et al., 2011). The relative climbing index was determined by splitting the climbing height into five equal parts and assigning scores from zero (at the base) to four (top section).

#### 2.9.5. Female fecundity and fertility assays

To assess the impact of ZnO-NPs treatment on female reproductive capacity, we performed controlled mating experiments using the w¹¹¹⁸line. Specifically, 15-day-old virgin females which had been maintained on a high-sugar diet (HSD) for 10 days with 5-day ZnO-NPs treatment were crossed 2-3-days-old young w¹¹¹⁸ virgin males maintained on a normal diet (ND).

Following mating at 25 °C, females were transferred to specialised egg-laying media consisting of 1% sucrose supplemented with yeast paste. Fecundity (the number of eggs laid) and fertility (the number of viable offspring) were then assessed (Kairamkonda & Upendra,2014).

#### 2.9.6. Male fertility assay

15-day-old virgin males,which had been maintained on a high-sugar diet (HSD), for 10 days followed by a 5-day ZnO-NPs treatment were crossed with 2-3-days old three young virgin females (1:3) maintained on a normal diet (ND), and then the male fertility was checked. Male fertility was quantified based on the number of viable F_1_ progeny emerging from each mating cross. The number of adult progeny was recorded after completion of development and expressed as progeny per mating cross. Reduced progeny production was considered indicative of impaired male reproductive performance (Chen P et al., 2021; Su et al., 2024) The fertility and morphological endpoints were evaluated comparatively among the ND control, HSD control and HSD + ZnO-NP treatment groups.

For cellular and spermatogenic assessment, dissected testes were fixed in 4% paraformaldehyde for 20-30 min at room temperature, followed by washing with PBS. Where applicable, nuclei were stained with DAPI and filamentous actin was visualized using fluorescent phalloidin to facilitate examination of the testicular architecture and developing spermatogenic cysts. Representative microscopic images were obtained from independent biological replicates. Testicular morphology was interpreted together with progeny production to determine whether changes in reproductive performance were accompanied by detectable alterations in testicular structure and spermatogenesis.

#### 2.9.7. RNA extraction, cDNA synthesis and RT-qPCR

Total RNA was isolated from healthy adult *D. melanogaster* using TRIzol reagent, following Singh et al. (2024) with minor modifications Flies were collected directly into chilled TRIzol, and three independent biological replicates were prepared for each group. RNA concentration and purity were determined spectrophotometrically, and integrity was verified by agarose gel electrophoresis. First-strand cDNA was synthesised using the Thermo First-Strand cDNA

Synthesis Kit according to the manufacturer’s protocol. Quantitative real-time PCR (RT-qPCR) was performed on a QuantStudio 3 Real-Time PCR System. The *rp*49 gene was used as the endogenous reference, and *Sod* was selected as the target gene. Primer sequences were adopted from published *D. melanogaster* RT-qPCR studies: *rp49* forward, 5′-GACGCTTCAAGGGACAGTATCTG-3′ and reverse, 5′-AAACGCGGTTCTGCATGAG-3′; *Sod* forward, 5′-TGGCCACATCAACCACAC-3′ and reverse, 5′-TTCCACTGCGACTCGATG-3′. Reactions were prepared using a SYBR Green master mix. The cycling protocol consisted of initial denaturation at 95°C for 12 min, followed by 40 cycles of denaturation at 95°C for 15s, annealing at 60°C for 20s, and extension at 72°C for 20s. Melt-curve analysis was performed to confirm product specificity. Ct values were recorded, and relative transcript abundance was calculated using the comparative 2^−ΔΔC^_t_ method, with *rp*49 as the reference gene and the untreated/control group as the calibrator. The 2^−ΔΔC^_t_ method was originally described by Livak and Schmittge (2001).

#### 2.9.8. Confocal imaging of brain and ovarian tissues

We dissected the ovaries of 14-day-old mated female progeny and brains from 14-day-old males and females from each treatment group in phosphate-buffered saline (PBS). The tissues were then processed for confocal imaging following the protocols described by Kairamkonda & Upendra (2014). High-resolution images were captured using the Dragonfly High-Speed Confocal Microscope System. For the subsequent qualitative and quantitative analysis, all images were processed and analyzed using Fiji/ImageJ (version 3.2.0.115).

## 3. Results and Discussion

The subsections below provide the results of the characterization of ZnO-NPs. Moreover, the anti-oxidant, anti-diabetic and anti-inflammatory potential on a high-sugar diet (HSD) using *Drosophila melanogaster* model.

### 3.1. Analytical Assays

#### 3.1.1. Plant-Based Extraction Mechanism for Synthesizing ZnO-NPs

The environmentally friendly method for producing ZnO-NPs utilizes plant phytochemicals that stabilize and cap compounds, facilitating the transformation of metal salts into metal nanoparticles. Important phytochemicals like alkaloids, aldehydes, terpenoids, flavonoids, and phenolic acids serve as stabilizers, inhibiting the clustering of particles. These phytocompounds, acting as reducing and capping agents, are present in diverse concentrations within various plant extracts. As a result, the composition of the plant fraction greatly influences the synthesis of nanoparticles. Zinc (II) ions found in plant extracts are reduced to metallic zinc, instead of creating a coordinated complex. Following the total reduction of the zinc precursor, metallic zinc interacts with dissolved oxygen in the solution, resulting in the creation of zinc oxide nuclei. The creation, stability, and generation of nanoparticles are affected by multiple factors, such as pH, concentration of metal salts, duration of contact, temperature, and the phytochemical composition of the plant fraction.

#### 3.1.2. Characterization of Biosynthesized Zinc Oxide Nanoparticles (ZnO-NPs)

The reflectance spectra of the prepared green ZnO-NPs using *F. macrophylla* is shown in Fig. 1. The absorption peak was observed at the wavelength 365 nm within the UV–Vis range of 300-500 nm. The reflectance peak at 365nm signifies the formation of pure, nanostructured wurtzite ZnO (Jamdagni et al., 2016)

**Fig. 1.**
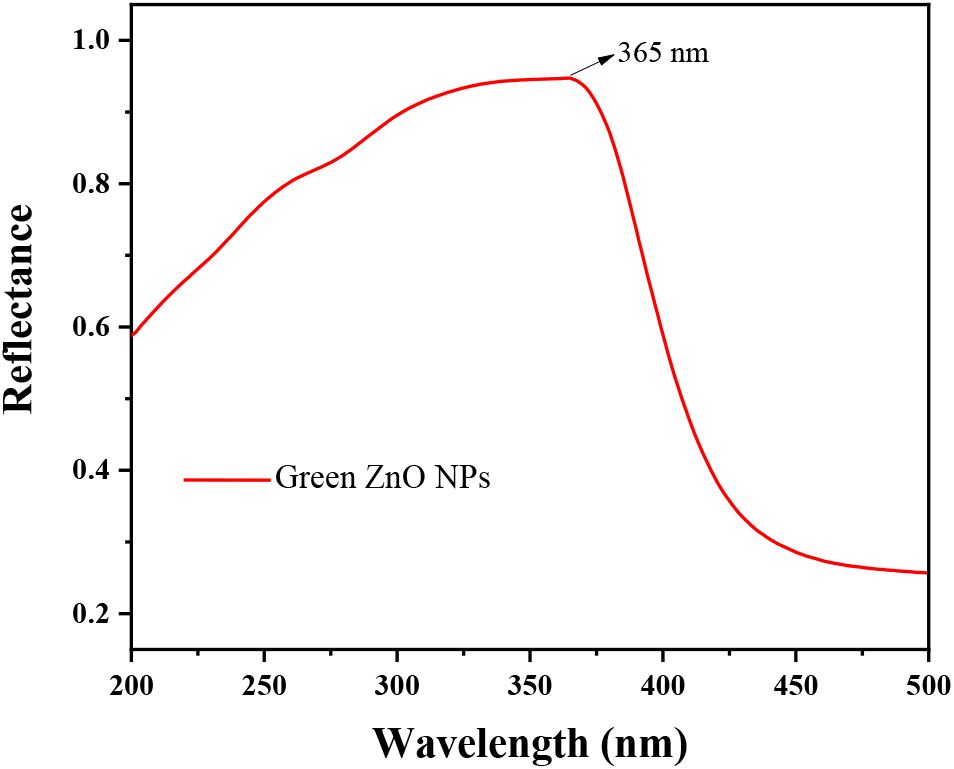
Reflectance spectra for ZnO-NPs

#### 3.1.3. Crystallographic Analysis of ZnO-NPs Using X-ray Diffraction

The crystallinity of green-synthesized ZnO-NPs was investigated using X-ray diffraction technique. Herein, the diffraction peak of the plane (100), (200), (101), (102), (110), (103), (200), (112), (201), (004) and (202) can be indexed to the 2θ value 31.97, 34.63, 36.45, 38.17, 47.73, 56.79, 6305, 66.56, 68.14, 69.25, 72.78 and 77.10 (shown in fig. 2) revealed the formation of hexagonal crystal system (reference JCPDS: 01-075-0576 with space group P63mc and space number 186). The crystallite size (D) of the prepared green ZnO-NPs (annealed at 500°C) is calculated using Debye Scherer’s equation from the prominent peak

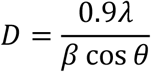

**Fig. 2.**
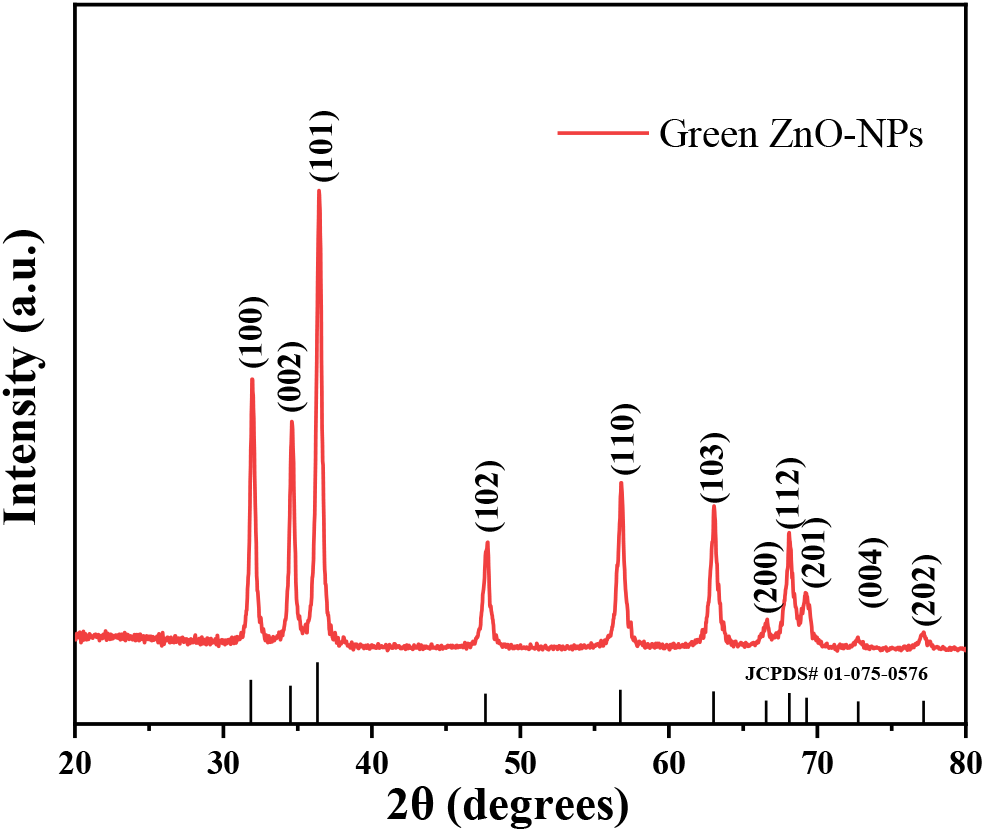
PXRD spectrum of Green ZnO-NPs

Where, λ=1.5404 Å with Cu Kα radiation, β= FWHM (full width at half maximum) of the dominant peak and D = crystallite size. The crystallite size of the (101) plane for green ZnO NPs is found to be 28.5 nm.

#### 3.1.4. Fourier Transform Infrared (FT-IR) Analysis

FT-IR spectra were employed within the wavelength range of 400–4000 cm^−1^ to identify the functional groups involved in the synthesis of ZnO-NPs derived from the aqueous extracts of FM leaves. The results from the FTIR analysis indicate that the surface of the nanoparticles is enriched with functional groups associated with phytoconstituents, including glycosides, alkaloids, tannins, reducing sugars, flavonoids, phenols, and others. This investigation revealed the presence of phenolic and hydroxyl groups, as evidenced by the broad energy bands observed between 3500 and 3200 cm^-1^, which are linked to the stretching vibrations of O-H groups. These groups are integral to the structures of lignin, cellulose, and hemicellulose. IR peak for as prepared (at 90°C):3365cm^-1^(O-H stretching) 1556 cm^-1^ (C=C/amine-NH stretching), 1392 cm^-1^ (C-H stretching), 1336cm^-1^ (C=O stretching), 1020 cm^-1^(C-O stretching), 422cm^-1^ (Zn-O). IR peak of ZnO-NPs (annealed at 500°C) 880cm^-1^ (ZnO), 420cm^-1^ (Zn-O) shown in fig. 3

**Fig. 3.**
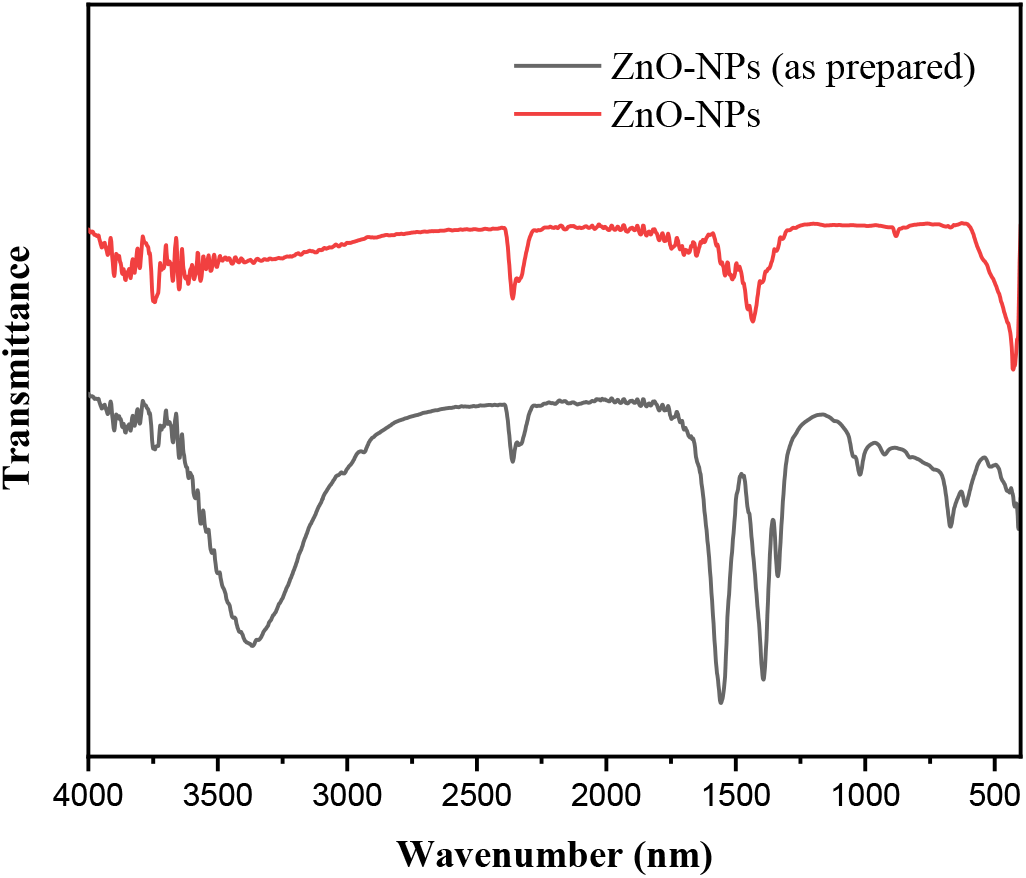
IR spectra of as prepared ZnO-NPs and annealed ZnO-NPs

#### 3.1.5. HRTEM analysis

HRTEM (High-resolution transmission electron microscope) analysis of *Flemingia macrophylla* mediated ZnO-NPs were conducted to calculate the morphological, particle size structural information shown in Fig.4(A-C), which visualized the green route of ZnO-NPs structural features. TEM micrograph of ZnO NPs shows quasi-spherical to irregular polyhedral nanoparticles, highlighting the structurally controlled morphology. The calculated average size of ZnO NPs calcined at 500°C is 47.13±9.33 nm (inset image shown in Fig. 4(A). HRTEM images of ZnO-NPs display the interplanar spacing of 0.28nm corresponding with hkl (100) shown in Fig. 4 (B). SAED patterns of ZnO NPs polycrystalline (shown in Fig. 4(C) with the d-spacing values (using ImageJ software) were in close agreement with the d-spacing values calculated from PXRD (reference JCPDS: 01-075-0576). This strong correlation confirms the crystalline nature and phase purity of *Flemingia macrophylla* mediated ZnO-NPs samples.

**Fig. 4.**
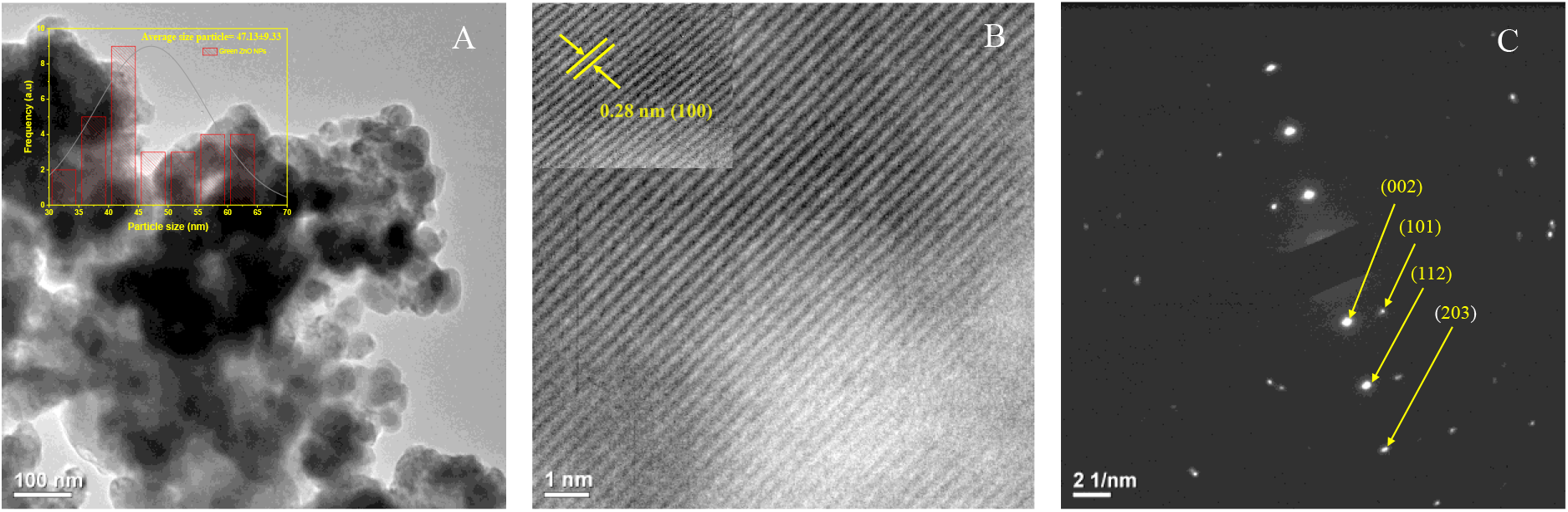
**(A)** TEM micrograph of ZnO NPs with histogram (inset) (**B**) HRTEM images of ZnO NPs. Fringes spacing for ZnO calculated using HRTEM image (inset) (**C**) SAED images of ZnO NPs

### 3.2. Determination of biological potential of the green synthesized ZnO-NPs

#### 3.2.1. Total phenolic content

The total phenolic content of the green-synthesized ZnO-NPs was determined using gallic acid as the reference standard. The gallic acid calibration curve showed a strong linear relationship between concentration and absorbance (y = 0.0068424x + 0.2746998; R² = 0.9974). Based on the absorbance of the ZnO-NP sample (0.5803), the phenolic content corresponded to 44.67 µg/mL gallic acid equivalents in the reaction mixture. After normalization to the amount of ZnO-NPs used in the assay, the total phenolic content was calculated as 446.68 mg gallic acid equivalents (GAE)/g sample.

#### 3.2.2. Total flavonoid content

The total flavonoid content (TFC) of the green-synthesized ZnO-NPs was determined using quercetin as the reference standard. The quercetin calibration curve showed a concentration-dependent increase in absorbance at 420 nm, with the regression equation y = 0.006722x + 0.15405. The ZnO-NPs sample (1 mg/mL) exhibited a mean absorbance of 0.4747, corresponding to 47.70 µg/mL quercetin equivalents based on the calibration curve. Accordingly, the TFC was 47.70 mg quercetin equivalents (QE)/g sample, assuming no additional dilution factor was applied. The result indicates the presence of appreciable flavonoid content in the green-synthesized ZnO-NPs.

### 3.3. Anti-oxidant activity

The anti-oxidant activity of green-synthesized ZnO-NPs was evaluated using DPPH, ABTS, and reducing-power assays (Fig. 5). ZnO-NPs showed a concentration-dependent increase in DPPH radical-scavenging activity, ranging from 5% at 10 µg/mL to 55% at 100 µg/mL, although the activity remained lower than that of ascorbic acid (IC_50_ = 95.06 ± 0.605 for ZnO-NPs and IC_50_= 18.67±1.29 for Ascorbic acid). A similar concentration-dependent pattern was observed in the ABTS assay, with inhibition increasing from 6% to 66% across 10–100 µg/mL with IC_50_=76.24 ± 1.49 but remaining below Trolox with IC_50_=33.985±1.07. The reducing power also increased progressively with ZnO-NPs concentration with an EC50 value of 37.90 ± 0.955 for ZnO-NPs compared to standard Ascorbic acid EC_50_=12.85± 0.688. Overall, ZnO-NPs demonstrated concentration-dependent radical-scavenging and reducing activities.

**Fig. 5.**
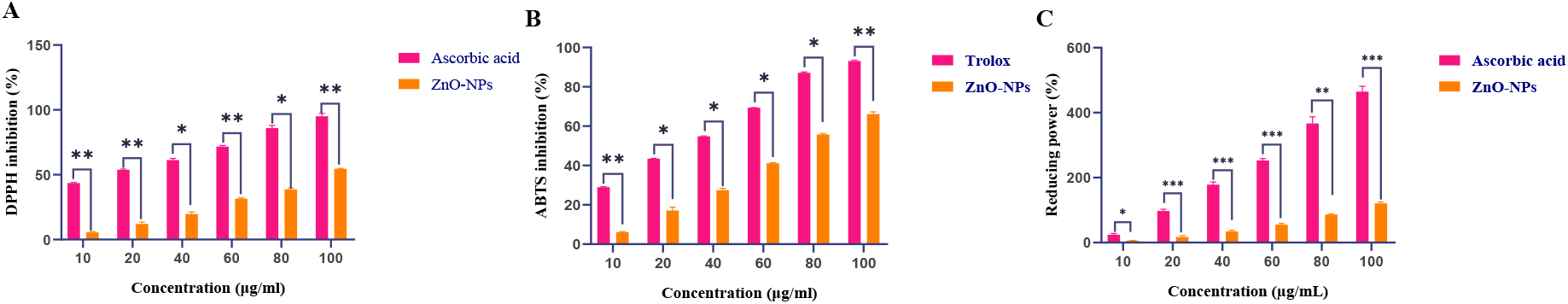
The graph depicts the antioxidant activity of Ascorbic acid/trolox and ZnO-NPs against (**A**)DPPH, (**B**)ABTS radical scavenging assay and (**C**) reducing power assay, respectively. Bars represent mean ± SD (** similar concentrations).

3.4. *In-vitro* Anti-diabetic activity:

ZnO-NPs exhibited concentration-dependent inhibitory activity against α-glucosidase, α-amylase, and lipase as shown in fig.6 (A,B and C). α-Glucosidase inhibition increased from approximately 6% at 10 μg/mL to 77% at 250 μg/mL, exceeding that of acarbose at 50-250 μg/mL with an IC_50_ value 159.19±1.199 µg/mL compared to 236±1.66 µg/mL of standard Acarbose. Similarly, α-amylase inhibition increased from 13% to 72% across the tested concentrations, approaching the inhibitory effect of Acarbose. The IC_50_ values were 126.56±1.509 µg/mL and 169.16±2.49 µg/mL respectively. ZnO-NPs also showed strong lipase inhibition (approximately 42-96%), exceeding the activity of orlistat at all tested concentrations with an IC_50_ of 72.31616±2.74 compared with 39.32±1.49 for standard

**Fig. 6.**
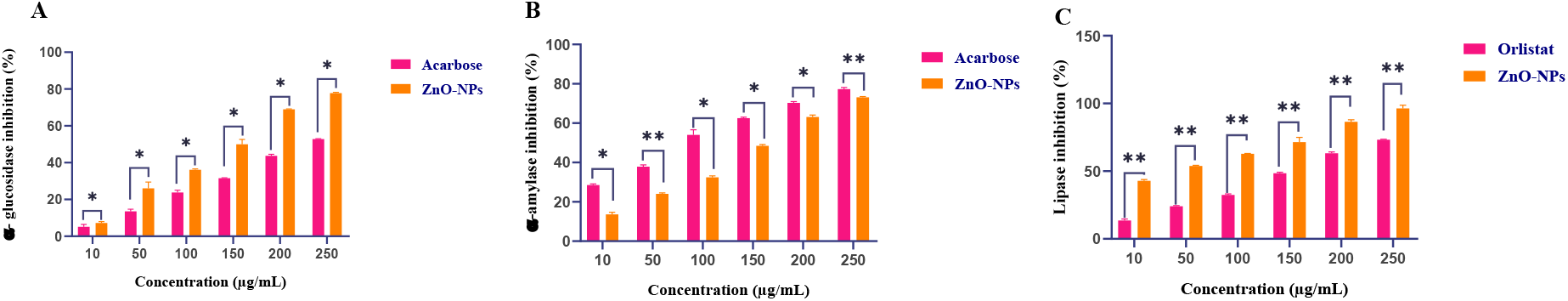
(**A**) α-glucosidase (**B**) α-Amylase inhibition and (**C**) Pancreatic lipase inhibitory effect of different concentrations of ZnO-NPs and Acarbose/orlistat as standards. Bars represent mean ± SD (ns **** p > 0.05, p < 0.0001 when compared at similar concentrations).

Orlistat. These findings indicate the potential of ZnO-NPs to inhibit key carbohydrate- and lipid-digesting enzymes, supporting their *in vitro* anti-diabetic activity.

### 3.5. Anti-inflammatory activity

The green-synthesized ZnO-NPs exhibited concentration-dependent anti-inflammatory activity in both assays as shown in Fig 7. In the protein denaturation assay, inhibition increased from 1.60% to 76.54% at 10–200 µg/mL, with an IC_50_ of 122.57 ± 1.16 µg/mL, compared with 104.88 ± 0.82 µg/mL for diclofenac. Similarly, ZnO-NPs inhibited heat-induced haemolysis from 17.63% to 75.91% at 10–250 µg/mL, with an IC_50_ of 153.17 ± 1.51 µg/mL, whereas indomethacin showed an IC_50_ of 105.12±1.65 µg/mL. These findings demonstrate concentration-dependent protein-stabilizing and membrane-protective activities of the ZnO-NPs, although their potency was lower than that of the reference drugs.

**Fig. 7.**
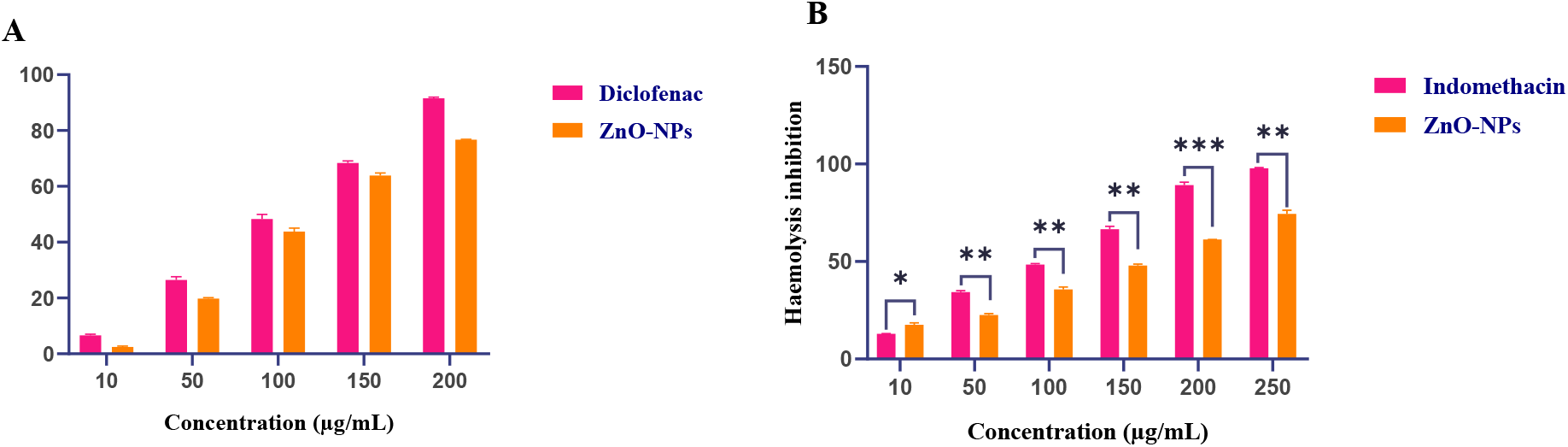
Invitro anti-inflammatory potential of ZnO-NPs and standard Diclofenac/Indomethacin extract by (**A**) protein denaturation inhibition activity (**B**) Heat induced Hemolysis

### 3.6. Cytotoxicity potential (MTT assay)

The green-synthesized ZnO-NPs were tested for toxicity against HeLa cells and normal HEK-293T cells by a cell viability assay. The results of the evaluation are demonstrated in Fig. 8.For HeLa cells, the cell viability significantly decreased with the enhancement of ZnO-NPs concentration if compared with the control group (about 90%). Specifically, the cell viabilities at 50, 100, 200, and 400 μM ZnO-NPs were around 87, 68, 55, and 40%, respectively shown in Fig. 8 (A). A marked decrease in cellviability indicates the cytotoxic activity of ZnO-NPs in a concentrationdependent manner. It can be estimated that the concentration of ZnO-NPs corresponding to 50% viability (IC_50_) was approximately 250-270 μM.

**Fig. 8.**
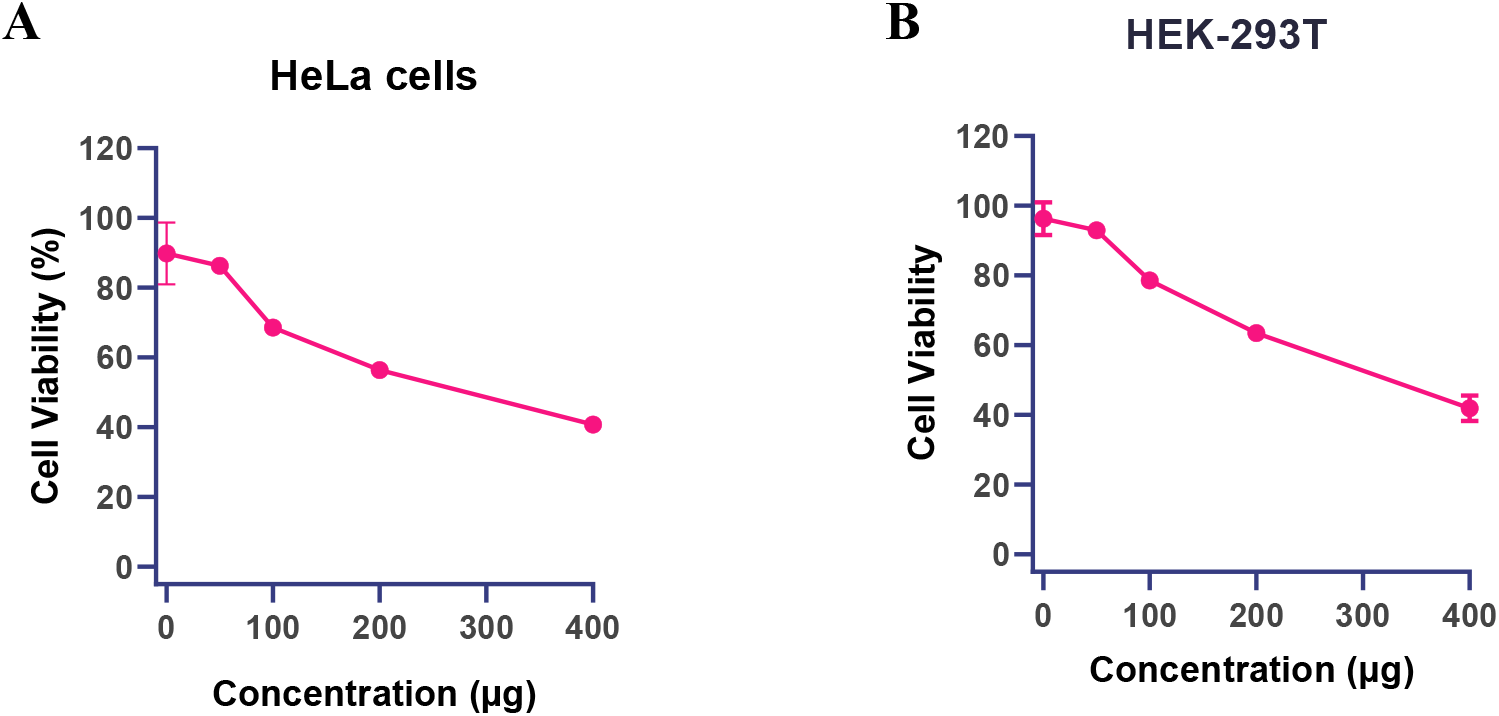
*In vitro* cytotoxicity profile of green-synthesized ZnO-NPs against cancerous **(A)** and non-cancerous cell lines **(B)**

For the treatment of non-cancerous HEK-293T cells, the similar tendency was also observed {Fig. 8 (B)}. The cell viability gradually declined with increasing concentration of ZnO-NPs from 96 to about 42% (for 0 and 400 μMZnO-NPs, respectively). The viability decreased to ∼50% at ∼300-320 μM ZnO-NPs.In the overall, the ZnO-NPs revealed dose-related toxic effect against both cancerous and normal cell lines.

3.7. *In-vivo* study on *Drosophila melanogaster* high-sugar-induced model:

#### 3.7.1. Estimation of glucose concentration in Drosophila

The HSD-induced *D. melanogaster* model exhibited elevated glucose levels, as indicated by increased glucose-associated absorbance. ZnO-NPs treatment progressively reduced absorbance from approximately 0.10 at 0.5 mg/10 g diet to 0.05 at 4 mg/10 g diet, with the 2 and 4 mg/10 g doses showing greater glucose-lowering effects than metformin under the experimental conditions. The glucose standard curve showed a concentration-dependent increase in absorbance over 25-300 μg/mL highlighted in Fig.9 and Fig.10 (A,B). Concordant trends obtained using GO-based and o-toluidine-based assays further supported the glucose-lowering effect of ZnO-NPs, particularly at higher doses.

**Fig. 9.**
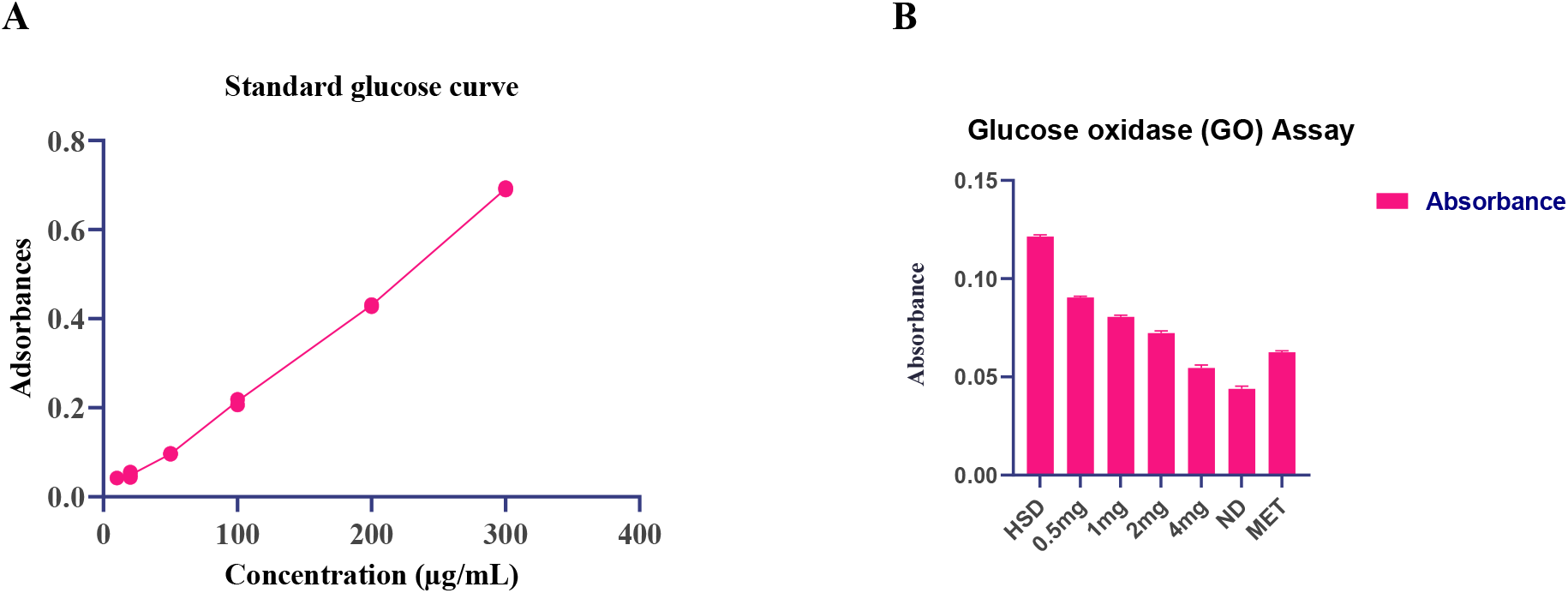
(**A**) Glucose standard curve for Glucose oxidase (GO) based colorimetric assay (**B**)Effect of ZnO-NPs on glucose levels in HSD-induced *D. melanogaster* compared with standard Metformin.

**Fig. 10.**
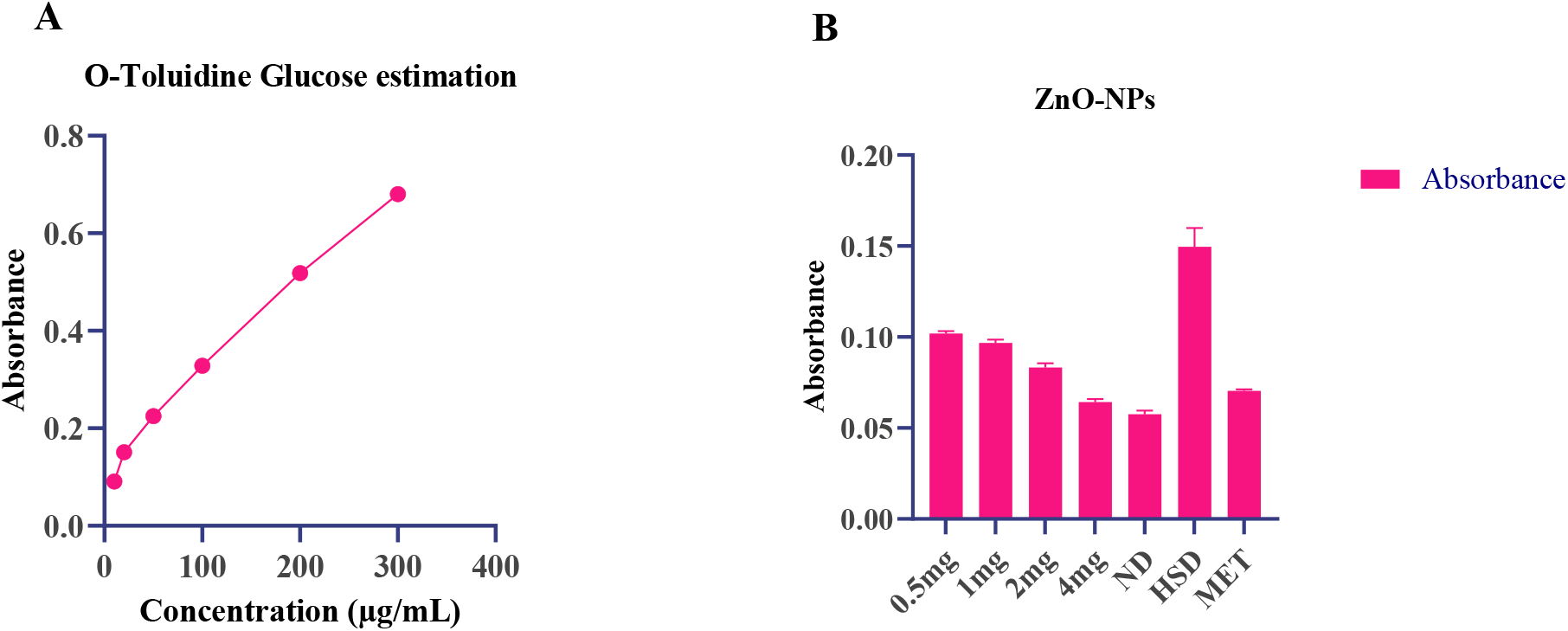
**(A)** Glucose standard curve for o-toluidine colorimetric method (**B**) Effect of ZnO-NPs on glucose levels in HSD-induced *D. melanogaster* compared with standard Metformin by o-toluidine colorimetric method.

Overall, the reduction in absorbance following ZnO-NPs treatment suggests a reduction in glucose levels in HSD-induced *D. melanogaster*, with greater reductions observed at the higher ZnO-NPs concentrations.

#### 3.7.2. *In-vivo* antioxidant assay

The *in-vivo* anti-oxidant activity of ZnO-NPs was evaluated using DPPH and ABTS radical-scavenging assays. Both assays demonstrated a concentration-dependent increase in anti-oxidant activity (Fig. 11A,B). DPPH inhibition increased from 49.07% at 0.5 mg/10 g diet to 76.88% at 4 mg/10 g, compared with 28.99% in the HSD group, with an IC_50_ of 0.54 mg/10 g diet. Similarly, ABTS inhibition increased from 44.73% to 76.40% across the same concentration range, compared with 28.87% in the HSD group, with an IC_50_ of 0.65 mg/10 g diet. These findings demonstrate a clear dose-dependent radical-scavenging activity of ZnO-NPs.

**Fig. 11.**
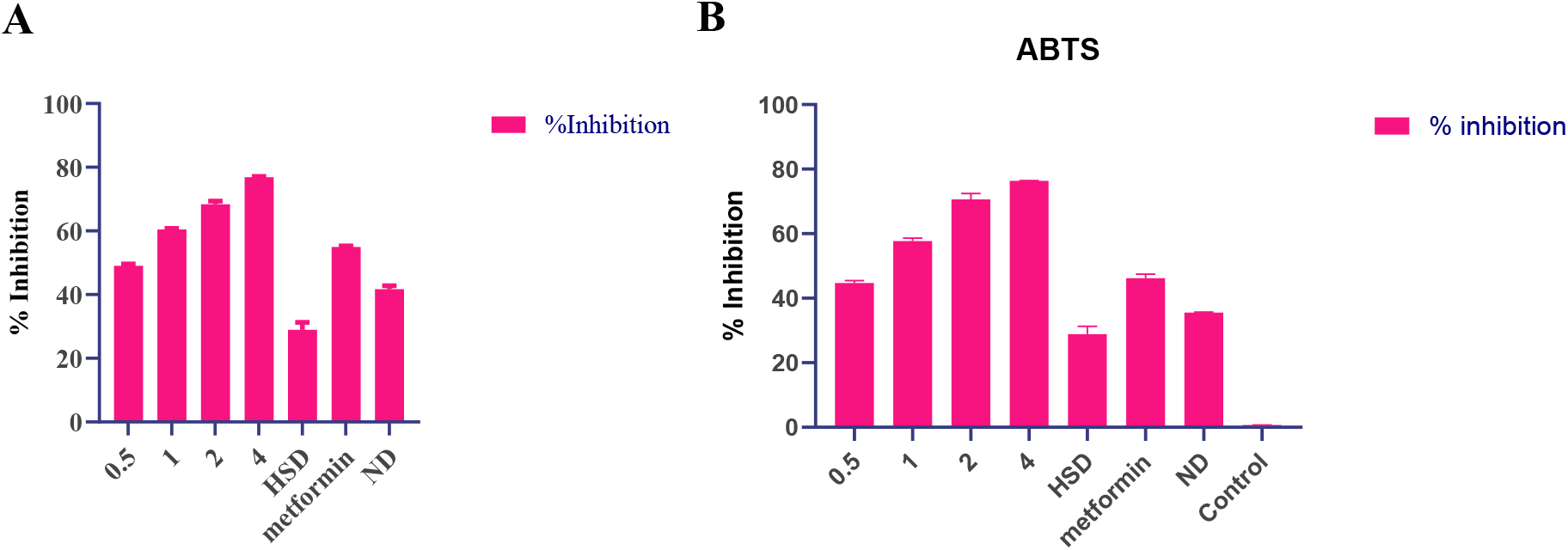
*In vivo* antioxidant activity of *F. macrophylla*-mediated ZnO-NPs in an HSD-induced *D. melanogaster* model (**A**) DPPH radical-scavenging activity and (**B**) ABTS radical-scavenging activity following treatment with ZnO-NPs at different concentrations and HSD, ND and Metformin standard.

#### 3.7.3. In-vivo anti-diabetic assay

The HSD group showed lower α-glucosidase inhibition (28.99%) than the normal diet group (37.01%). ZnO-NPs treatment produced a concentration-dependent increase in α-glucosidase inhibition, from 48.43% at 0.5 mg/10 g diet to 75.10% at 4 mg/10 g diet (Fig. 12). The highest dose showed an inhibitory activity comparable to metformin (77.25% at 16 mg/10 g diet), with an IC₅₀ of 0.4343 mg/10 g diet. These findings demonstrate a dose-dependent α-glucosidase inhibitory effect of ZnO-NPs in HSD-induced *D. melanogaster*.

**Fig. 12.**
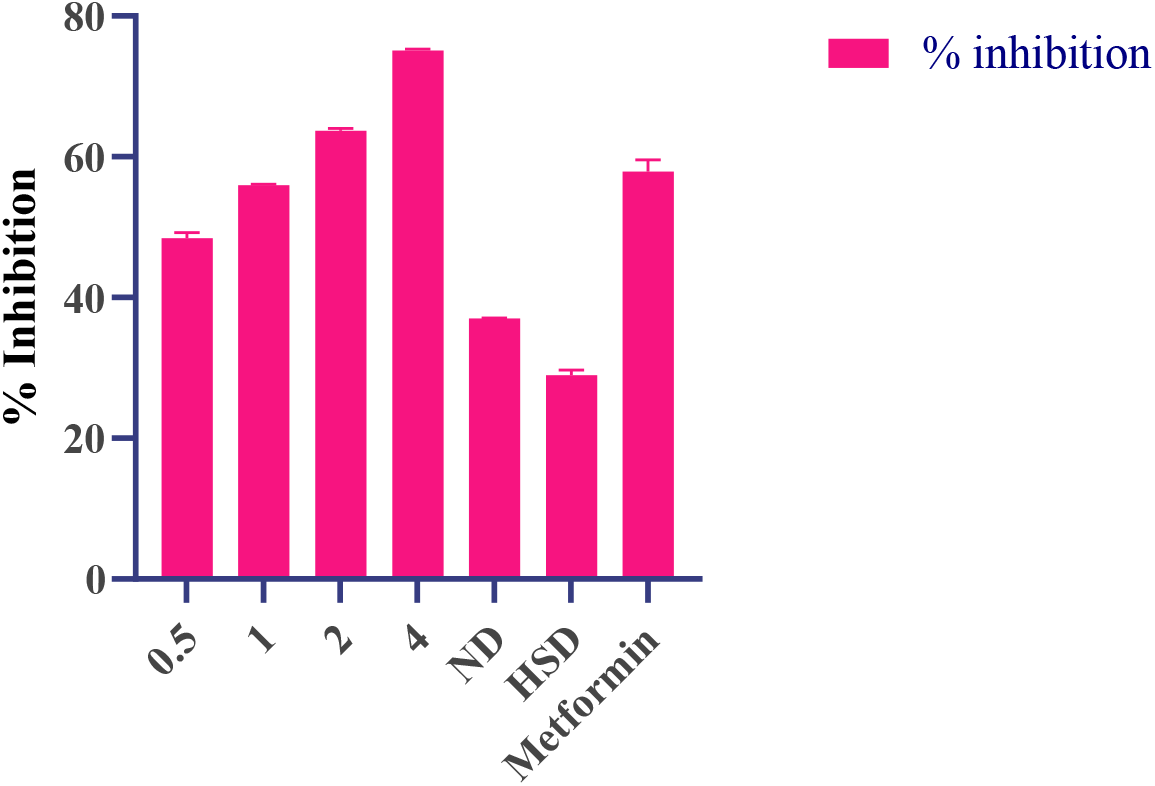
*In vivo* α-glucosidase inhibitory activity of ZnO-NPs in HSD-induced *D. melanogaster*.

Overall, the treatment showed strong activity against the enzyme to lower the level of blood glucose in the model under study. Specifically, increasing the concentration of the treatment improved its ability to suppress α-glucosidase, thereby reducing the rate of saccharide digestion and glucose absorption.

#### 3.7.4. Green-Synthesized ZnO-NPs Extended Lifespan and Modulated Stress-Responsive Gene Expression of Sod in High-Sugar Diet-Fed Drosophila

To evaluate the systemic therapeutic efficacy of green-synthesized ZnO-NPs, we first monitored adult longevity in both male {Fig. 13 (A)} and female {Fig. 13 (B)} *D. melanogaster* subjected to a High-Sugar Diet (HSD). Chronic exposure to HSD significantly compromised survival parameters compared to Normal Diet (ND) controls. Supplementation with phytogenic ZnO-NPs across varying concentrations (0.5, 1, 2, and 4 mg/mL) markedly improved lifespan profiles in a dose-dependent manner. Notably, the higher dosage treatments (2 mg/mL and 4 mg/mL) conferred a robust survival benefit that was comparable to, or exceeded, the positive therapeutic control Metformin (Met) (16mg/10gdiet).

**Fig. 13.**
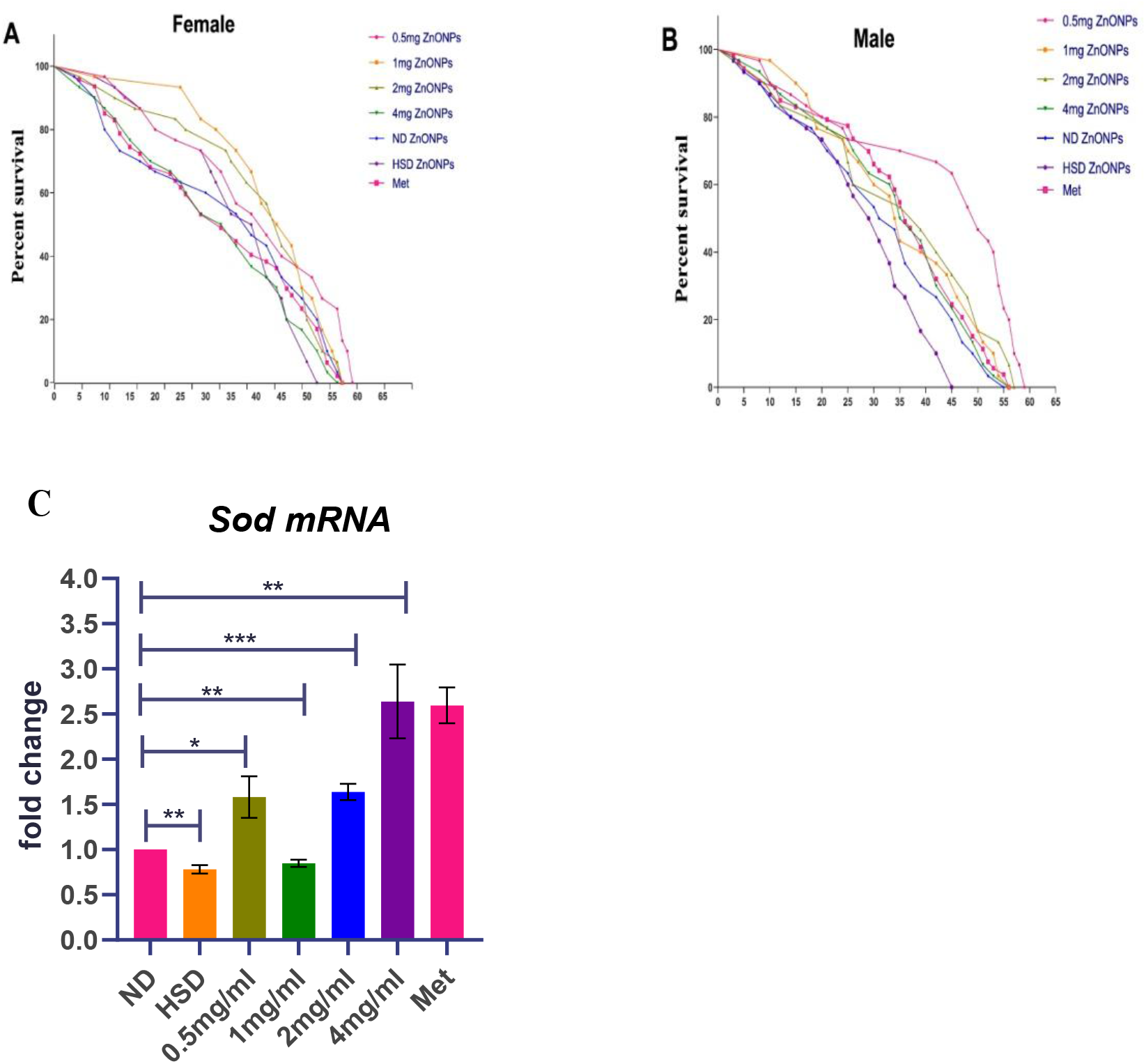
Effect of green-synthesized ZnO-NPs on longevity and transcript expression of *Sod* in *D. melanogaster*. (**A**–**B**) Kaplan-Meier survival curves showing percent survival over time (days) in female (A) and male (B) flies across experimental groups: Normal Diet (ND), High-Sugar Diet (HSD), HSD supplemented with varying concentrations of phytogenic ZnO-NPs (0.5, 1, 2, and 4 mg/mL), and Metformin (Met)(16mg/10g diet) as a positive control. (**C**) Real-time quantitative PCR (RT-qPCR) transcript fold-change analysis of endogenous antioxidant enzyme *Superoxide Dismutase* (*Sod*). Data are presented as mean ±SD (∗ *p* < 0.05,∗∗ *p* < 0.01, ∗∗∗ *p* < 0.001, ns = not significant).

To elucidate the molecular mechanisms underlying this survival advantage, we performed real-time quantitative PCR (RT-qPCR) to quantify transcript levels of the endogenous antioxidant enzyme *Superoxide Dismutase* (*Sod*).Flies maintained on HSD exhibited a significant reduction in *Sod* mRNA expression compared to ND controls {Fig.13 (C)}, *p* < 0.01), reflecting severe HSD-induced oxidative stress. Co-treatment with ZnO-NPs effectively rescued and upregulated *Sod* transcription in a dose-dependent fashion, with the 4 mg/mL treatment demonstrating a profound ≈ 2.5-fold increase relative to ND control (*p* < 0.01), equivalent to the response observed in Metformin-treated flies.

#### 3.7.4. ZnO-NPs Treatment mitigates HSD-induced vacuole formation in *Drosophila* brain and Neuromotor impairment in a dose-dependent manner

To evaluate the neuroprotective potential of green-synthesized zinc oxide nanoparticles (ZnO-NPs) under high-sugar conditions, we examined adult *Drosophila* central brain microarchitecture using Phalloidin (F-actin, red) and Hoechst (nuclei, blue) fluorescence staining {Fig. 14 (A)}. Brains from High-Sugar Diet (HSD)-fed flies displayed marked structural deterioration and increased tissue vacuolation compared to Normal Diet (ND) controls. Quantitative assessment of neuropathology at 10 days post-treatment revealed a significant increase in the average number of neuropil vacuoles in the HSD group relative to ND flies{Fig.14(B)}, *p* < 0.05. Co-treatment with phytogenic ZnO-NPs effectively preserved brain structural integrity and attenuated vacuole formation in a concentration-dependent manner. Specifically, flies treated with 2 mg/mL and 4 mg/mL ZnO-NPs exhibited a significant reduction in brain vacuoles (*p* < 0.05 relative to HSD), returning neuropathological counts to levels comparable to ND controls and the positive control Metformin.

**Fig. 14.**
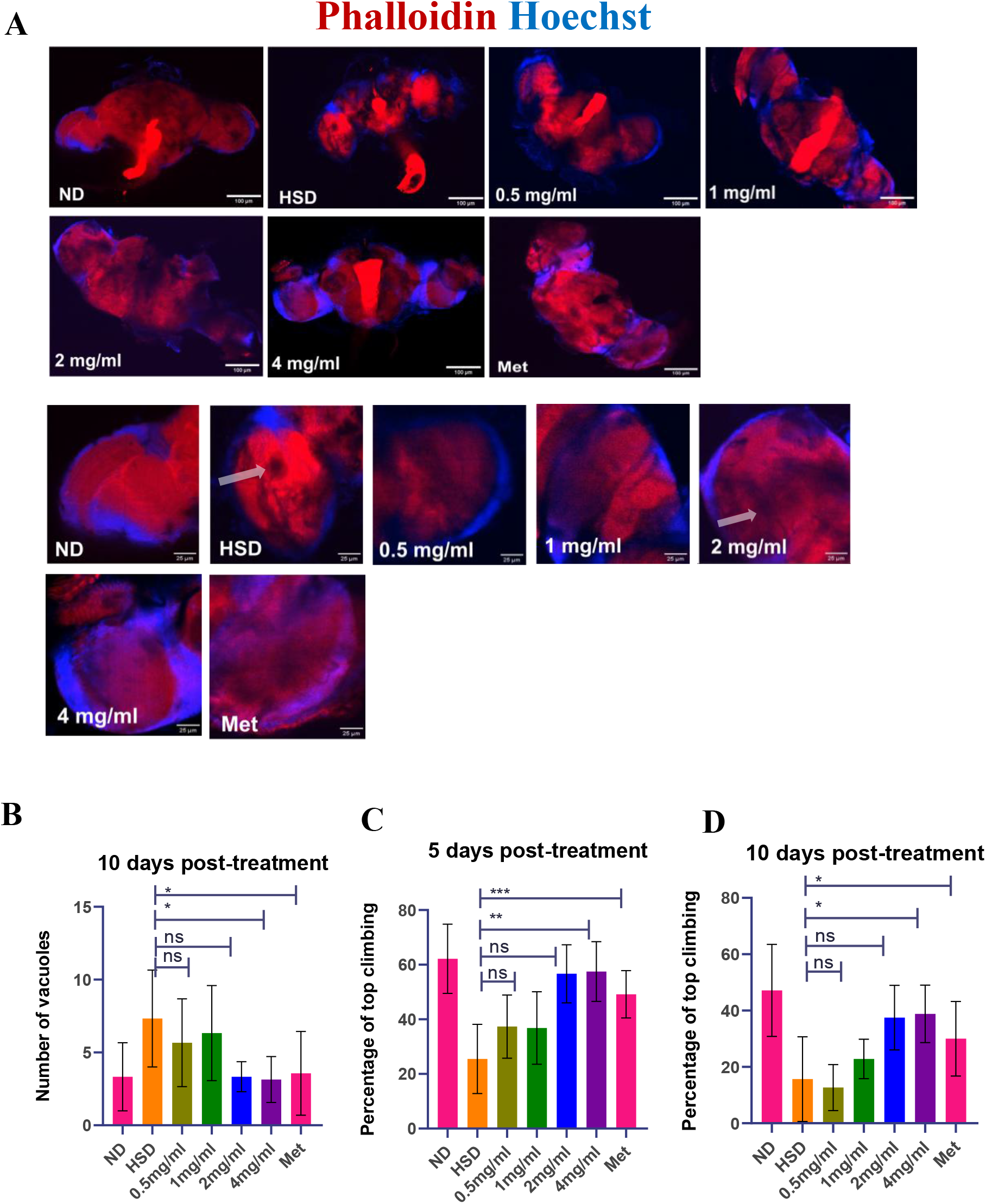
ZnO-NPs preserve brain neuroanatomical integrity, reduce neurodegeneration, and restore motor function in HSD-fed *D. melanogaster*. (**A**) Representative confocal fluorescent microphotographs of adult brains stained with Phalloidin (F-actin, red) and Hoechst (nuclei, blue) across treatment groups: Normal Diet (ND), High-Sugar Diet (HSD), HSD co-treated with green ZnO-NPs (0.5, 1, 2, and 4 mg/mL), and Metformin (Met) (16mg/10g) diet as a positive control. Lower panels show magnified views of optic lobe/brain microarchitecture. Scale bars = 100 *μ*m (whole brain) and 25 *μ*m (insets). (**B**) Quantitative estimation of brain neurodegeneration expressed as the number of vacuoles per brain at 10 days post-treatment, showing an HSD-induced increase in neurodegenerative vacuolation and its significant attenuation upon higher ZnO-NPs doses (2 and 4 mg/mL) and Metformin. **(C–D)** Rapid iterative negative geotaxis (climbing assay) displaying the percentage of flies reaching the top mark at **(C)** 5 days post-treatment and **(D)** 10 days post-treatment. ZnO-NPs supplementation (2 and 4 mg/mL) significantly rescues HSD-induced locomotor impairment. Data represent mean ±SD (∗ *p* < 0.05,∗∗ *p* < 0.01, ∗∗∗ *p* < 0.001, ns = not significant).

To determine whether the structural preservation of central brain microarchitecture correlated with functional motor recovery, we evaluated climbing performance using a rapid iterative negative geotaxis assay at 5 days {Fig. 14 (C)} and 10 days {Fig. 14 (D)} post-treatment. Chronic HSD consumption induced progressive motor impairment, resulting in a dramatic decrease in the percentage of flies reaching the top of the vial compared to ND controls (*p* < 0.001 at day 5; *p* < 0.05 at day 10). Supplementation with ZnO-NPs restored climbing ability in a dose- and time-dependent manner. While lower concentrations (0.5 mg/mL and 1 mg/mL) produced modest improvements, higher concentrations (2 mg/mL and 4 mg/mL) significantly rescued motor deficits at both 5 days (*p* < 0.01 and *p* < 0.001, respectively) and 10 days (*p* < 0.05) post-treatment. Notably, the locomotor performance in the 2 mg/mL and 4 mg/mL ZnO-NPs treatment groups was statistically indistinguishable from ND controls. Taken together, these findings demonstrate that green ZnO-NPs protect central nervous system architecture against HSD-induced neurotoxicity and prevent secondary locomotor decline.

#### 3.7.6. Green-Synthesized ZnO-NPs Ameliorate HSD-Induced Female Reproductive Dysfunction and Ovarian Atrophy

To determine the impact of green-synthesized ZnO-NPs on reproductive performance under metabolic stress, we evaluated fecundity, fertility, and ovarian morphology in female *D. melanogaster*(Fig.15). Chronic exposure to HSD severely impaired female reproductive capacity, causing a sharp decline in egg-laying activity {Fig. 15 (A)} and hatching efficiency

**Fig. 15.**
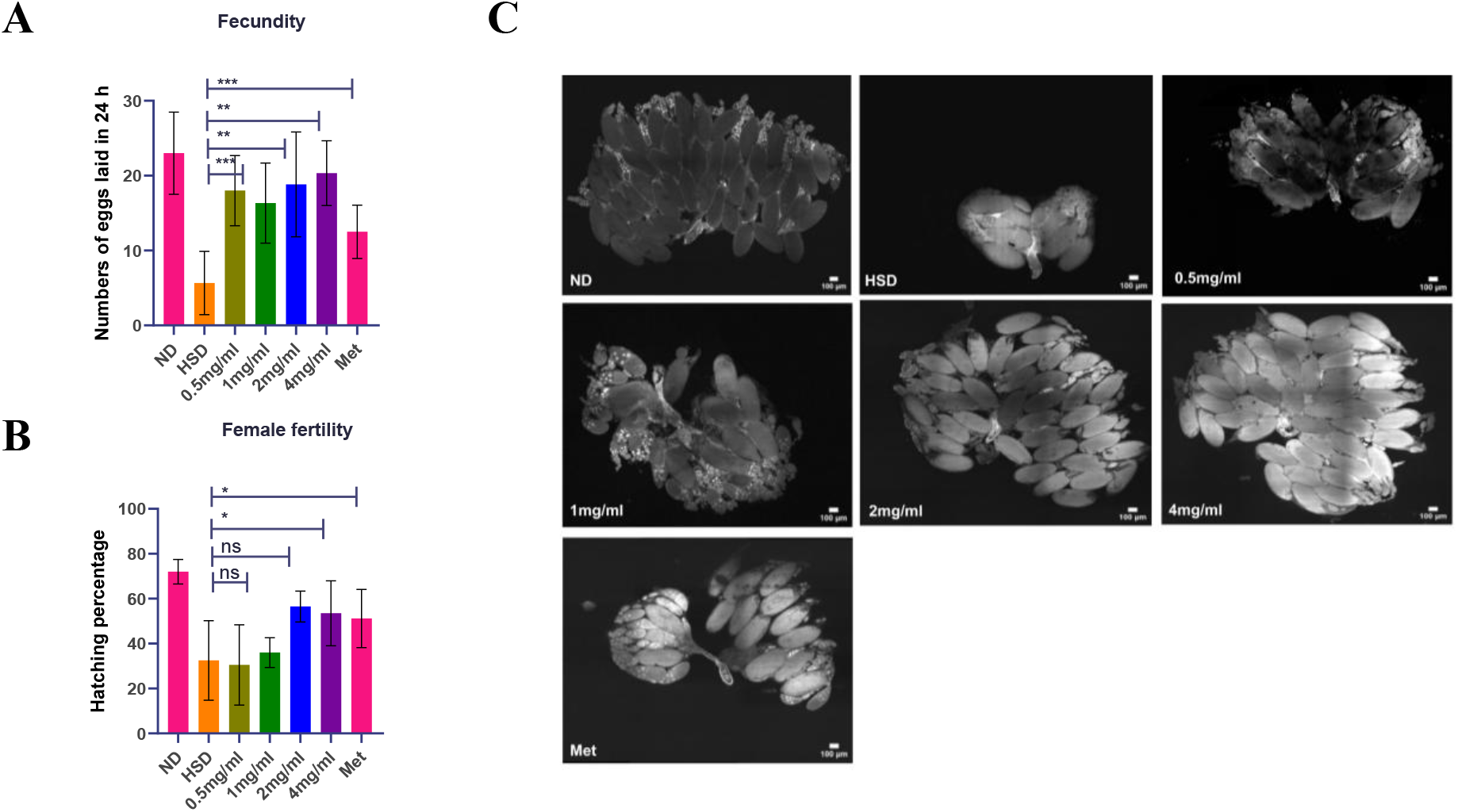
Impact of green ZnO-NPs on female fecundity, fertility, and ovarian morphology. (**A**) 24-hour egg production, (**B**) hatching percentage, and (**C**) representative microphotographs of dissected ovaries across experimental groups (ND, HSD, 0.5-4.0 mg/10g diet ZnO-NPs, and Metformin 16mg/10g diet). Green ZnO-NPs dose-dependently mitigate HSD-induced reproductive suppression and ovarian tissue atrophy (∗ *p* < 0.05,∗∗ *p* < 0.01, ∗∗∗ *p* < 0.001, ns = not significant). Scale bars = 100 *μ*m.

{Fig. 15(B)} compared to ND controls. Supplementation with phytogenic ZnO-NPs (0.5, 1, 2, and 4 mg/mL) substantially rescued egg production in a dose-dependent manner. All ZnO-NPs treatment concentrations (0.5-4 mg/mL) significantly restored fecundity relative to the HSD group (*p* < 0.01 and *p* < 0.001), elevating oviposition rates to levels comparable to the standard antidiabetic control Metformin (Met).

Similarly, evaluation of female fertility demonstrated that HSD feeding dramatically reduced embryonic hatching success (≈ 30% hatching rate). While lower ZnO-NPs concentrations (0.5 and 1 mg/mL) did not significantly alter hatching efficiency compared to HSD (*p* = ns), higher concentrations (2 and 4 mg/mL) significantly rescued hatching success (*p* < 0.05), increasing hatching percentages to near-control and Metformin levels {Fig. 15 (B)}.

To confirm whether functional reproductive recovery was associated with structural integrity, we examined adult ovary morphology across treatment groups {Fig. 15 (C)}. Dissected ovaries from HSD-fed females exhibited severe structural degeneration, characterized by pronounced ovarian atrophy, underdeveloped oocytes, and reduced egg chambers relative to the fully developed ovaries of ND flies. Co-treatment with green ZnO-NPs mitigated HSD-induced gonadal damage in a concentration-dependent manner, with 2 mg/mL and 4 mg/mL doses restoring mature egg chamber abundance, oocyte morphology, and overall organ size to levels matching ND and Metformin controls. These findings demonstrate that green ZnO-NPs protect female gonadal microarchitecture and mitigate HSD-induced reproductive decline.

#### 3.7.7. Assessment of Male Fertility and Testicular Microarchitecture Following ZnO- NPs Supplementation

To determine whether HSD-induced metabolic stress affects male reproductive output and whether green-synthesized zinc oxide nanoparticles (ZnO-NPs) confer protective effects, we evaluated male fertility and male reproductive tract morphology (Fig. 16). Quantitation of male fertility, expressed as the number of adult progeny eclosed, revealed a slight reduction in offspring number in the HSD-fed group compared to Normal Diet (ND) controls

**Fig. 16.**
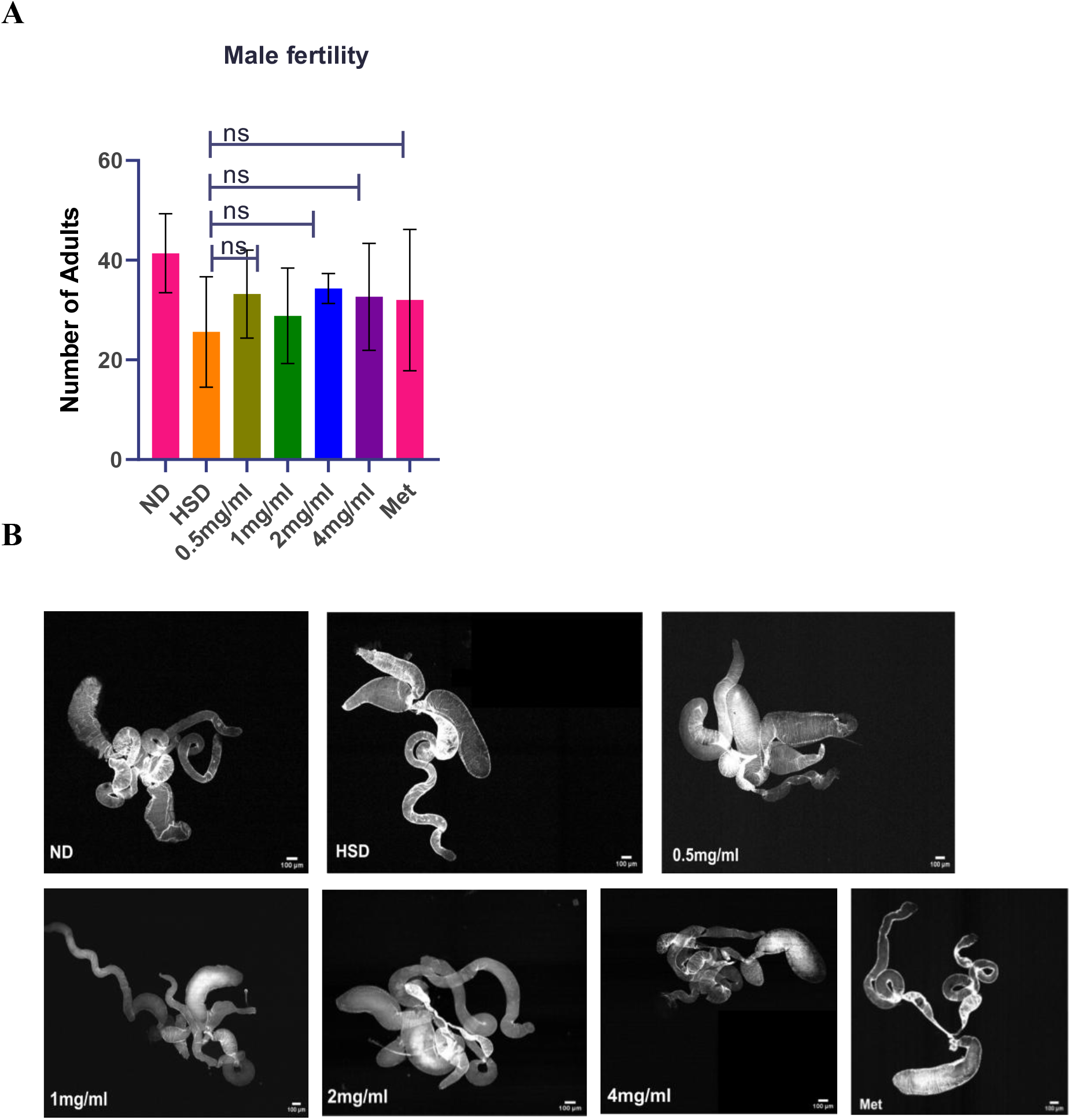
Effect of green-synthesized ZnO-NPs on male reproductive parameters and organ integrity. (**A**) Male fertility expressed as the total number of adult progenies eclosed per male. Differences across ND, HSD, ZnO-NPs (0.5, 1, 2, and 4 mg/mL), and Metformin (Met) 16mg/10g diet groups remain statistically non-significant (ns). **(B)** Representative light micrographs of dissected adult male reproductive tracts demonstrating preserved testis, seminal vesicle, and accessory gland morphology across all treatment conditions. Scale bars = 100 *μ*m.

{Fig.16(A)}. Co-treatment with green ZnO-NPs (0.5, 1, 2, and 4 mg/mL) and Metformin (Met) showed a trend toward restoring adult progeny yield closer to ND baseline levels; however, these differences across groups were statistically non-significant (*p* = ns).

Microscopic examination of dissected adult male reproductive tractscomprising the testes, seminal vesicles, and accessory glandsrevealed intact tubular architecture and morphology across all experimental conditions {Fig.16(B)}. Testes from HSD-fed males displayed no severe structural damage or gross atrophy compared to ND controls. Supplementation with phytogenic ZnO-NPs (0.5-4.0 mg/10g diet) maintained normal organ morphology, structural integrity, and tubular coiled conformation comparable to both ND and Metformin-treated controls, indicating that green ZnO-NPs exhibit no gonadotoxic effects on the male reproductive system at these concentrations.

### 3.8. Discussion

The present study demonstrates the multifunctional biological effects of F. macrophylla in the preparation of zinc oxide nanoparticles (ZnO-NPs), which reduce metabolic disorder, oxidative stress, and neurological and reproductive disturbances caused by a high-sugar diet (HSD) in D. melanogaster. The crystalline (hexagonal wurtzite) ZnO-NPs with crystallite size of an average particle size 47.13 ± 9.33 nm were successfully synthesised in accordance with earlier reports of the plant-mediated synthesis of ZnO NPs, where phytochemicals are responsible for reducing, nucleating, stabilising, and functionalizing the precursor (Issam et al., 2021; Thema et al., 2015; Aldeen et al.,2022; Naseer et al.,2020) [2,5,11,18]. The presence of detectable phenolic and flavonoid compounds suggests that metabolites of F. macrophylla remain bound to the nanoparticle surface, highlighting their potential role in bioactivity, given the reported antioxidant and anti-inflammatory properties of Flemingia species (Gahlot et al.,2013; Kumbhare & Ukey, 2023). The concentration-dependent antioxidant activity observed in DPPH, ABTS, and reducing assays suggests potential redox-modulating activity, likely due to synergistic interactions between the ZnO surface and phytochemical components. Botanically relevant because too much sugar in the diet disrupts redox homeostasis and causes oxidative damage. HSD-fed flies showed decreased antioxidant levels and Sod expression, with improved responses to ZnO-NP treatments. However, the restoration of Sod transcription plays a special role in the detoxification of superoxide, a finding that awaits confirmation at the protein and enzymatic levels. *In vitro* enzyme assay provided mechanistic insights into the inhibitory effects on α-glucosidase, α-amylase, and pancreatic lipase, confirming interference with enzymes involved in the digestion of carbohydrates and lipids. The distinct α-glucosidase inhibition is significant, as inhibition of intestinal α-glucosidase will minimise the generation and absorption of glucose from the intestine (Kumar et al., 2011). Neuropraxia ZnO-NPs inhibited glucose-associated responses consistently, with a higher effect noted when a larger number of ZnO-NPs was fed to HSD-fed flies. Glucose reduction is, however, not only due to α·glucosidase inhibition, because the existence of the effects of zinc on biological activity suggests that ZnO can also modulate metabolic cell processes, oxidative homeostasis and insulin-like signalling (Jiang et al., 2018; Tang, 2019). This improvement in survival speaks to the benefits observed at the organism level in mitigating metabolic and oxidative stress. The neurobehavioral dysfunction caused by HSD was also improved. The lack of brain vacuolation was observed, and brain returns negative axes were observed in ZnO-NPs at higher concentration. The metabolic protection and neurobehavioral improvement occurred simultaneously, and their combination suggests a possible relationship between metabolic protection mechanisms and neurological benefits.

However, the specific neuroprotective pathways have yet to be determined, and future investigations should focus on elucidating neuronal apoptosis, mitochondrial activity, ROS accumulation, and insulin/FOXO signalling to better elucidate the mechanisms. A unique result was the restoration of the female reproductive function. High HSD levels reduced fecundity, hatching efficiency, and OV morphology, which were alleviated by supplementation with ZnO-NPs. This is consistent with the previously observed sensitivity of Drosophila reproductive fitness to nutritional imbalance and oxidative stress [36]. The impact was found to be sex-dependent; however, as male fertility production was not affected, there was no alteration in the morphology of the testicles and accessory glands, whereas in the females, there was a morphological alteration. Importantly, concentration-dependent cytotoxicity in HeLa and HEK 293T cells indicates that the bioactivity of the ZnO-NPs is dose-dependent and that “green synthesis” does not always mean the product will be completely safe. These types of problems have also been addressed in previous biomedical research (Jiang et al., 2018; Mishra et al., 2017). Therefore, higher doses resulted in greater antioxidant and metabolic responses, but these need to be taken in conjunction with cytotoxicity observations, indicating the need for extensive mammalian safety studies. Overall, the consistent responses observed across all biochemical assays, glucose regulation, antioxidant defence, survival, Sod expression, neuroanatomical integrity, locomotor behaviour, and female reproductive functions suggest that F. macrophylla mediated ZnO-NPs exert several beneficial effects against HSD-induced metabolic stress. Rather, the findings are considered in vitro and largely rely on a Drosophila model, and thus must be viewed as preclinical proof of concept and should not be interpreted as proof of therapeutic efficacy. Only after future studies define the characteristics of the phytochemical corona and release of Zn²⁺, pin down the contribution of plant-derived compounds to ZnO, and elucidate insulin/metabolic and inflammatory signalling pathways, as well as mammalian safety profiles, can the translational potential be determined.

### 3.9. Conclusions

The present study established an integrated green-synthesis and biological evaluation platform for *F. macrophylla*-mediated ZnO-NPs. The aqueous leaf extract successfully supported the formation of crystalline ZnO-NPs exhibiting characteristic optical, structural, and morphological properties, including a hexagonal wurtzite phase with an average particle size of 47.13±9.33 nm following calcination.

The synthesized ZnO-NPs demonstrated concentration-dependent antioxidant, carbohydrate-hydrolyzing enzyme inhibitory, pancreatic lipase inhibitory, and membrane/protein-stabilizing activities *in vitro*. In the HSD-induced *D. melanogaster* model, ZnO-NPs supplementation reduced glucose-associated responses, enhanced *in vivo* antioxidant capacity, increased α-glucosidase inhibitory activity, improved survival, restored *Sod* expression, attenuated HSD-associated brain vacuolation, and improved locomotor performance. Female reproductive dysfunction was also substantially alleviated, as reflected by improved fecundity, hatching efficiency, and ovarian morphology. In males, ZnO-NPs treatment did not significantly alter fertility and was associated with preservation of testicular and accessory reproductive tract morphology under the conditions tested.

Collectively, these findings indicate that *F. macrophylla*-mediated ZnO-NPs possess multifunctional biological activity against several phenotypic consequences of high sugar dietary stress, with antioxidant and metabolic effects potentially contributing to the observed neuroprotective and reproductive benefits. The restoration of *Sod* expression provides additional evidence that modulation of endogenous antioxidant defenses may participate in the protective response.

At the same time, the concentration-dependent cytotoxicity observed in HeLa and HEK-293T cells emphasizes the importance of dose optimization and comprehensive safety assessment. Therefore, the present findings delineate preclinical proof-of-concept evidence in a *Drosophila* model rather than evidence of established therapeutic efficacy. Future studies should define the nanoparticle surface chemistry, clarify the contribution of *F. macrophylla* phytochemicals versus ZnO-NPs itself, investigate insulin/metabolic and inflammatory signalling pathways, and evaluate pharmacokinetics, biodistribution, genotoxicity and mammalian efficacy. Such studies will be essential to determine whether the promising multifunctional activity of these phytogenic ZnO-NPs can be translated into a safe and mechanistically defined nanotherapeutic strategy for metabolic disease.

## Data availability

The data supporting the findings of this study are available within the paper. Any additional data connected to the study are available from the corresponding author upon reasonable request.

## CRediT authorship contribution statement

**Lhaineichong Khongsai-** Writing – original draft, Validation, Methodology, Investigation, Formal analysis, Data curation, Conceptualization. **Rounab Sarkar-** Writing, Visualization, Software, Methodology, Data curation.: Validation, Conceptualization. **Atom Rajiv Singh-** Writing, Visualization, Validation, Methodology, Formal analysis. **Asem Sanjit Singh-** Review & editing. **BK Bindhani** - Supervision, Conceptualization. **Upendra N-** Review & editing. **Ch. Brajakishor Singh**- Review & editing, Supervision, Funding acquisition, Conceptualization

## Declaration of competing interest

The authors declare that they have no known competing financial interests that could have appeared to influence the work reported in this paper.

## Acknowledgements

Authors are grateful to the Director, IBSD for financial support and providing infrastructures to carry out the experiment. Authors are thankful to Developmental Biology and Genetics (DBG) lab, IISc. Bengaluru for providing infrastructures and Drosophila model to conduct the experiments. The authors also gratefully acknowledge the Instrumentation Facility, North-Eastern Hill University (NEHU), for providing the necessary instrumentation facilities for the characterization of the biosynthesized zinc oxide nanoparticles (ZnO NPs)

## Abbreviations

ABTS: 2,2′-azino-bis(3-ethylbenzothiazoline-6-sulfonic acid)
DPPH: 2,2-diphenyl-1-picrylhydrazyl
FT-IR: Fourier transform infrared spectroscopy
HSD: high-sugar diet
HRBC: human red blood cell
ND: normal diet
NPs: nanoparticles
PXRD: powder X-ray diffraction
RING: Rapid Iterative Negative Geotaxis
RT-qPCR: real-time quantitative PCR
SAED: selected area electron diffraction
SEM: scanning electron microscopy
TEM: transmission electron microscopy
UV–Vis: ultraviolet-visible spectroscopy
ZnO-NPs: zinc oxide nanoparticles.

## Notes

### Competing Interest Statement

The authors have declared no competing interest.

